# The root-knot nematode effector Mj-MSP18: a Swiss army knife for reprogramming plant immunity

**DOI:** 10.1101/2025.08.28.672664

**Authors:** Yujin Chen, Muhammad I. Maulana, Jana Oklestkova, Jitka Široká, Veronique Jonckheere, Hugues De Gernier, Lander Bauters, Ning Zhang, Miroslav Strnad, Petra Van Damme, Alain Goossens, Godelieve Gheysen, Boris Stojilković

## Abstract

Root-knot nematodes (*Meloidogyne* spp.) are obligatory plant root parasites whose effector proteins play a critical role in suppressing plant immunity. However, the effectors direct host targets and underlying molecular mechanisms remain poorly understood. Using TurboID-mediated proximity labeling in tomato (*Solanum lycopersicum*) hairy roots, we identified an interaction between the nematode effector Mj-MSP18 and the tomato BRASSINOSTEROID-SIGNALING KINASE 7 (Sl-BSK7). Yeast two-hybrid (Y2H) assays confirmed that this interaction is conserved in *Arabidopsis thaliana* and *Nicotiana benthamiana*. Additionally, yeast three-hybrid and luciferase complementation assays demonstrated that Mj-MSP18 disrupts the interaction between BSK7/8 and FLS2 in both yeast and *in planta*. Given that the BSK7/8–FLS2 interaction is essential for flg22-induced pattern-triggered immunity (PTI), this disruption likely accounts for the suppression of reactive oxygen species (ROS) production and callose deposition observed upon transient expression of Mj-MSP18 in flg22-treated *N. benthamiana* leaves. Correspondingly, *Arabidopsis* and tomato *bsk7* mutants exhibited increased susceptibility to root-knot nematode infection.

Furthermore, RNA-seq analysis of tomato hairy roots expressing *Mj-MSP18* revealed extensive transcriptional reprogramming, including the downregulation of defence-related genes and hydrogen peroxide response pathways. In addition, Y2H screening identified Sl-MYB and Sl-MYC2 as additional interactors, linking Mj-MSP18 to phytohormone biosynthesis, particularly the brassinosteroid (BR) and salicylic acid (SA) pathways, as validated by targeted metabolite analysis. The conservation of Mj-MSP18 across *Meloidogyne* species suggests a broadly conserved mechanism for host immune suppression and phytohormone modulation.

## Introduction

Root-knot nematodes (RKNs, *Meloidogyne* spp.) are obligate phytopathogens that infect a wide range of important crops, including tomato (1), and are ranked among the most invasive plant disease-causing agents worldwide (2). Infective vermiform second-stage juveniles (J2s), which hatch from eggs in the soil, are attracted to root tip. They penetrate the root elongation zone and migrate intercellularly to reach the plant vascular cylinder, where they establish specialized, multinucleate feeding structures known as giant cells to extract water and nutrients during their sedentary biotrophic lifestyle (3). Giant cell formation results from successive nuclear divisions without cytokinesis, followed by isotropic cell expansion (4). Simultaneously, surrounding cells divide and undergo vascular differentiation, leading to the formation of characteristic root knots, or galls. Over several weeks, the nematodes intermittently feed from various giant cells and progress through successive molts to become adult females, which produce hundreds of eggs deposited onto the root surface in a protective gelatinous matrix called the egg mass (5).

To penetrate roots and establish feeding sites, RKNs utilize a specialized, needle-like stylet that mechanically pierces plant cell walls and allows access to intracellular contents. The stylet also serves as a conduit for the secretion of numerous effector proteins into the apoplast or directly into host cells. These effectors facilitate root invasion, suppress plant immunity, and regulate giant cell development (6, 7). For example, the *Meloidogyne* effector MiMsp40 has been shown to suppress PAMP elf18-triggered immunity and Bax-induced cell death in plants (8). More recently, OPR2 (12-oxophytodienoate reductase 2), a key enzyme in JA biosynthesis, was found to be targeted by *M. incognita* effector MiMSP32 to suppress host defense (9).

From a host perspective, to counteract pathogen invasion, plants have evolved various defense strategies. As the first layer of the plant immune system, pathogen-associated molecular patterns (PAMPs) or damage-associated molecular patterns (DAMPs) released by RKNs are perceived by cell-surface-localised pattern recognition receptors (PRRs), initiating pattern-triggered immunity (PTI) during host invasion (10, 11). For instance, the ascaroside Ascr#18 from nematodes (12) is perceived by NEMATODE-INDUCED LRR-RLK1 (NILR1) to trigger PTI (13). PTI is accompanied by a series of downstream immune responses, including the generation of reactive oxygen species (ROS), activation of mitogen-associated protein kinases (MAPKs), deposition of callose at the cell wall, and host transcriptome reprogramming, all of which contribute to pathogen defense (14, 15).

Plant PRRs primarily consist of receptor-like kinases (RLKs) and receptor-like proteins (RLPs), which enable early detection and broad-spectrum resistance against pathogens (16). Both RLKs and RLPs contain extracellular domains responsible for the PAMP/DAMP perception and a transmembrane domain, but only RLKs possess a cytoplasmic kinase domain. Additionally, plants encode numerous receptor-like cytoplasmic kinases (RLCKs), which are evolutionarily related to RLKs but lack both extracellular and transmembrane domains (17). Leucine-rich repeat (LRR) receptor-like kinases are the largest group of RLKs in plants. A well-characterized example is FLAGELLIN-SENSING 2 (FLS2), which recognizes the flg22 epitope of bacterial flagellin to activate immune responses (18, 19). Upon flg22 perception, FLS2 forms a complex with BRASSINOSTEROID INSENSITIVE 1 (BRI1)-ASSOCIATED RECEPTOR KINASE 1 (BAK1), initiating downstream signaling such as ROS production and MAPK activation (20–22). In addition, recent studies have revealed that certain *Arabidopsis* RLCKs, known as BRASSINOSTEROID-SIGNALING KINASES (BSKs), are associated with FLS2 and are required for flg22-induced PTI against bacteria and fungi (23–25).

In this study, we investigated the molecular mechanism by which the *M. javanica* effector Mj-MSP18 suppresses plant immunity. *Mi-MSP18*, a homolog of *Mj-MSP18*, was previously shown to be upregulated during nematode parasitic stages, with overexpression in rice enhancing RKN parasitism and RNAi silencing reducing it (26). Using TurboID-mediated proximity labeling, yeast two-hybrid assays, and confocal microscopy, we identified an interaction between Mj-MSP18 and a tomato receptor-like cytoplasmic kinase homologous to *Arabidopsis* BSK7 (Sl-BSK7), which is associated with Sl-FLS2 and involved in flagellin-triggered PTI (27). Quantitative yeast three-hybrid and luciferase complementation assays elucidated that Mj-MSP18 can disrupt the BSK7-FLS2 interaction. Additionally, Sl-BSK7 plays a role in plant defense against RKNs. Through Y2H screening, we identified tomato MYB and MYC2 transcription factors as additional interactors of Mj-MSP18, which may explain the effector-induced transcriptional reprogramming observed in the host. Finally, targeted metabolite analysis supported the role of Mj-MSP18 in modulating phytohormone pathways. Altogether, our findings identify key host targets of Mj-MSP18 and propose a molecular mechanism by which nematodes suppress plant immunity, offering new insights into plant-RKN interactions.

## Results

### Mj-MSP18 is a conserved *Meloidogyne* effector

To gain deeper insights into the molecular mechanisms of effectors involved in the *M*. *javanica*–*Solanum lycopersicum* interaction, we investigated *M. incognita* effectors and their closest homolog from *M. javanica* (28–31). One of them, *M. incognita* putative oesophagus gland cell secretory protein 18 (NCBI accession number: AAN08584) (32), was shown to be upregulated during parasitic stages, and its overexpression in rice enhances RKN parasitism (26). We investigated the closest *M. javanica* homolog, which shares 98.3% protein identity with the *M. incognita* version (**Figure S1A**). *Mj-MSP18* is predicted to encode a protein of 172 amino acids with a molecular mass of 18.6 kDa. The N-terminal 20-amino acids are predicted to form a signal peptide for secretion from the dorsal gland into the plant (Huang et al., 2003). The version of the effector protein lacking this signal is referred to as Mj-MSP18_ΔSP_.

Mj-MSP18 is a conserved effector across various *Meloidogyne* species, including *M. incognita, M. hapla*, *M. arenaria*, *M. enterolobii* and *M. graminicola,* as evidenced by the phylogenetic tree generated with Clustal Omega **(Figure S1B)** using protein databases from these RKN species (*Meloidogyne* INRAE and WormBase ParaSite). This high degree of conservation across species suggests an evolutionary important role during infection.

### Proximity labeling identifies plant proximal partners of the nematode effector Mj-MSP18

We used TurboID-mediated proximity labeling (33–35) to identify proximal plant protein targets (i.e., proxeome) of the nematode effector Mj-MSP18 in tomato hairy roots. Mj-MSP18 localizes to both the nucleus and cytoplasm (**Figure S2).** The TurboID tag was C-terminally fused to Mj-MSP18_ΔSP_ and eGFP, and these fusion proteins were expressed in tomato hairy roots under the control of an estrogen-inducible promoter as described previously (34). Both fusion proteins were well expressed and biotinylated, as confirmed by anti-FLAG and streptavidin blotting, except for the streptavidin-bound fraction of Mj-MSP18_ΔSP_ **(Figure S3)**. This discrepancy may be attributed to the previously reported influence of biotinylation on epitope tag detection, especially for lysine-containing tags (e.g., FLAG)— where masking effects can impair detection despite adequate protein expression and biotinylation (36). Biotinylated proteins captured from high-quality, reproducible samples **(Figure S4)** were analyzed by LC-MS (37, 38). Several candidate proteins were significantly enriched in the Mj-MSP18_ΔSP_ samples compared to the control, as highlighted in the volcano plot **(Figure 1A, Supplemental Table 1)**. Among these, Sl-BSK7 (*Solyc12g099830*), a receptor-like cytoplasmic kinase from *S. lycopersicum*, was prioritized for further analysis due to its strong enrichment—a 16-fold increase in abundance—and statistical significance (FDR ≤ 0.05; p-value ≤ 0.001) in the Mj-MSP18_ΔSP_ TurboID dataset, as well as its known role in immune signaling.

**Figure 1.**
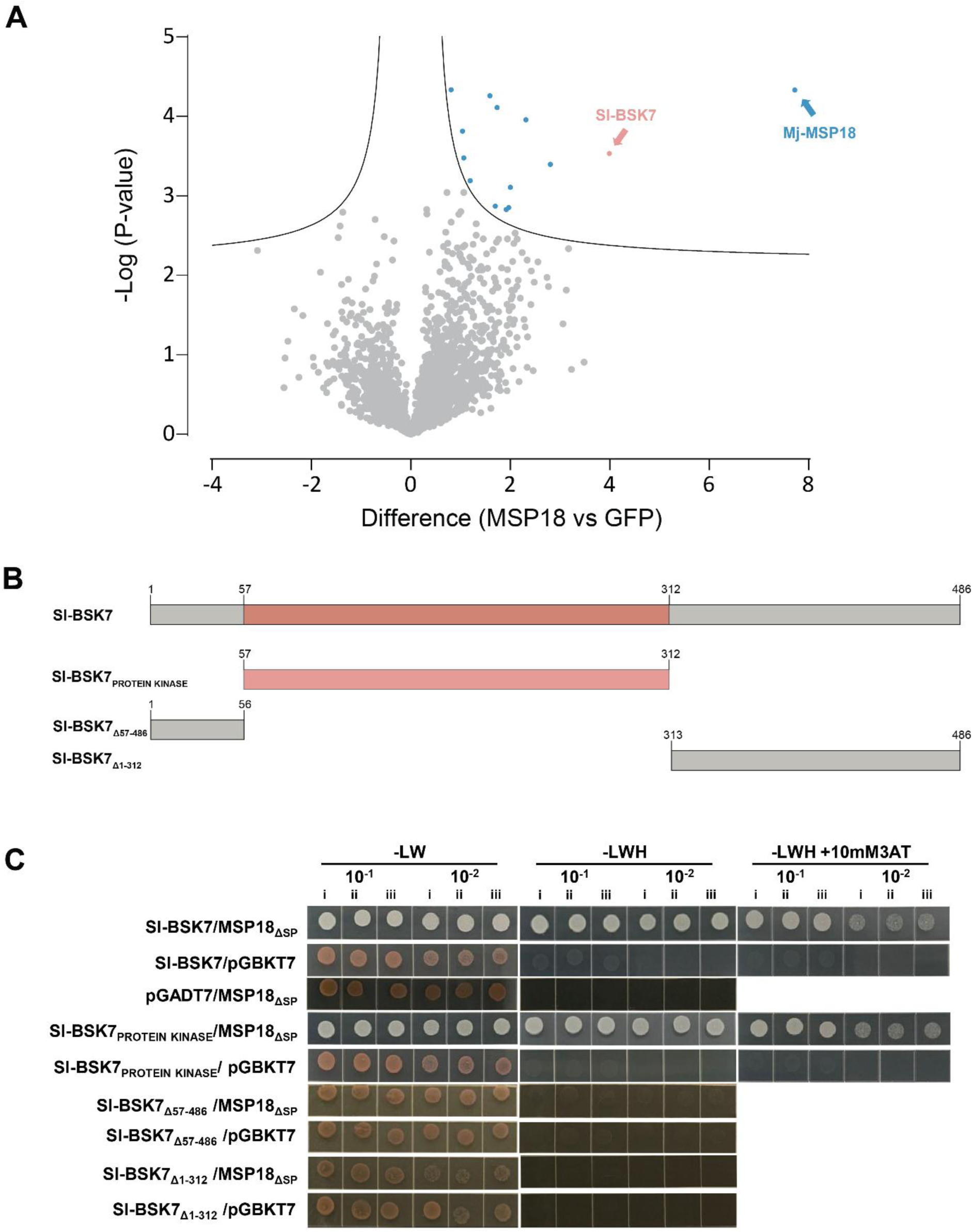
Turbo-ID and Y2H reveal interaction between Mj-MSP18_ΔSP_ and Sl-BSK7. **(A)** Volcano plot showing pairwise comparisons of Turbo-ID results between Mj-MSP18_ΔSP_ and GFP. A two-sample *t*-test was used to identify significantly enriched plant proteins in the Mj-MSP18_ΔSP_ Turbo-ID samples compared to the GFP control, using a false discovery rate (FDR) of 0.05 with S0 = 0.1 (i.e. artificial within groups variance; this value defines the relative importance of the P value and difference between means). The *t*-test difference is plotted on the x-axis, and –log(*P*-value) on the y-axis. Proteins enriched in Mj-MSP18_ΔSP_ are on the right, and depleted proteins on the left. Significantly enriched hits are marked in blue; Sl-BSK7 is highlighted in pink. **(B)** Schematic representation of the tomato SL-BSK7 protein, showing the protein kinase domain and its truncated versions. **(C)** Y2H analysis of the interaction between Mj-MSP18_ΔSP_ and various Sl-BSK7 truncated constructs. PJ69-4A yeast cells were co-transformed with Mj-MSP18_ΔSP_ and each truncated version of Sl-BSK7. Transformants were plated on SD (Synthetic Defined) medium lacking leucine and tryptophan (-LW) or additionally histidine (-LWH), with or without 3-amino-1,2,4-triazole (3-AT) to test for interaction strength. Autoactivation controls consisted of the same yeast strain co-transformed with each Sl-BSK7 construct and the empty vector. Truncated protein versions are indicated on the left; dilution series are indicated on the top. i, ii and iii indicate independent biological replicates.

### Y2H validates direct interaction between MSP18_ΔSP_ and Sl-BSK7

We performed a yeast two-hybrid (Y2H) analysis with Mj-MSP18_ΔSP_ as the bait, confirming that Sl-BSK7, a plant immunity-related proximal partner from the TurboID proxeome list, can directly interact with Mj-MSP18 **(Figure 1C, Supplemental Table 1)**. Sl-BSK7 was also cloned into several truncated versions, encompassing the N-terminal part (Sl-BSK7_Δ57-486_), the C-terminal part (Sl-BSK7_Δ1-312_), and the isolated protein kinase domain (Sl-BSK7_PROTEIN KINASE_). The results showed that Mj-MSP18_ΔSP_ specifically recognizes and strongly binds to Sl-BSK7_PROTEIN KINASE_. Conversely, no direct interactions were detected with other parts of the Sl-BSK7 protein **(Figure 1C)**, indicating that the protein kinase domain is essential and sufficient for the interaction.

Homologous proteins of Sl-BSK7 from *A. thaliana*, *N. benthamiana*, and *S. lycopersicum* were identified based on sequence similarity with the protein kinase domain, and a phylogenetic tree and sequence alignments were generated (**Figure S5, S6A**). BSK7 (AT1G63500) and BSK8 (AT5G41260) from *A. thaliana* (At-BSK7 and At-BSK8), as well as Nb-BSK7 (Niben101Scf04732c01004) from *N. benthamiana*, were tested for interaction with Mj-MSP18_ΔSP_. Strong interactions were observed with BSK proteins from all species tested **(Figure S6B)**.

Multiple sequence alignment of Sl-BKS7 and two *N. benthamiana* homologs (Niben101Scf04732g01004 and Niben101Scf 08430g00010) revealed high sequence conservation in the C-terminal region of the proteins (**Figure 2A**). Additional validation of Mj-MSP18_ΔSP_ binding was conducted using a smaller catalytic domain of the Sl-BSK7 protein kinase domain (Sl-BSK7_SER-THR/TYR-PROTEIN KINASE_), as well as the relatively conserved C-terminal regions of both tobacco orthologs. This analysis further narrowed down the critical interaction region in Sl-BSK7 to Sl-BSK7_Δ1-149Δ309-486_ **(Figure 2B)**.

**Figure 2.**
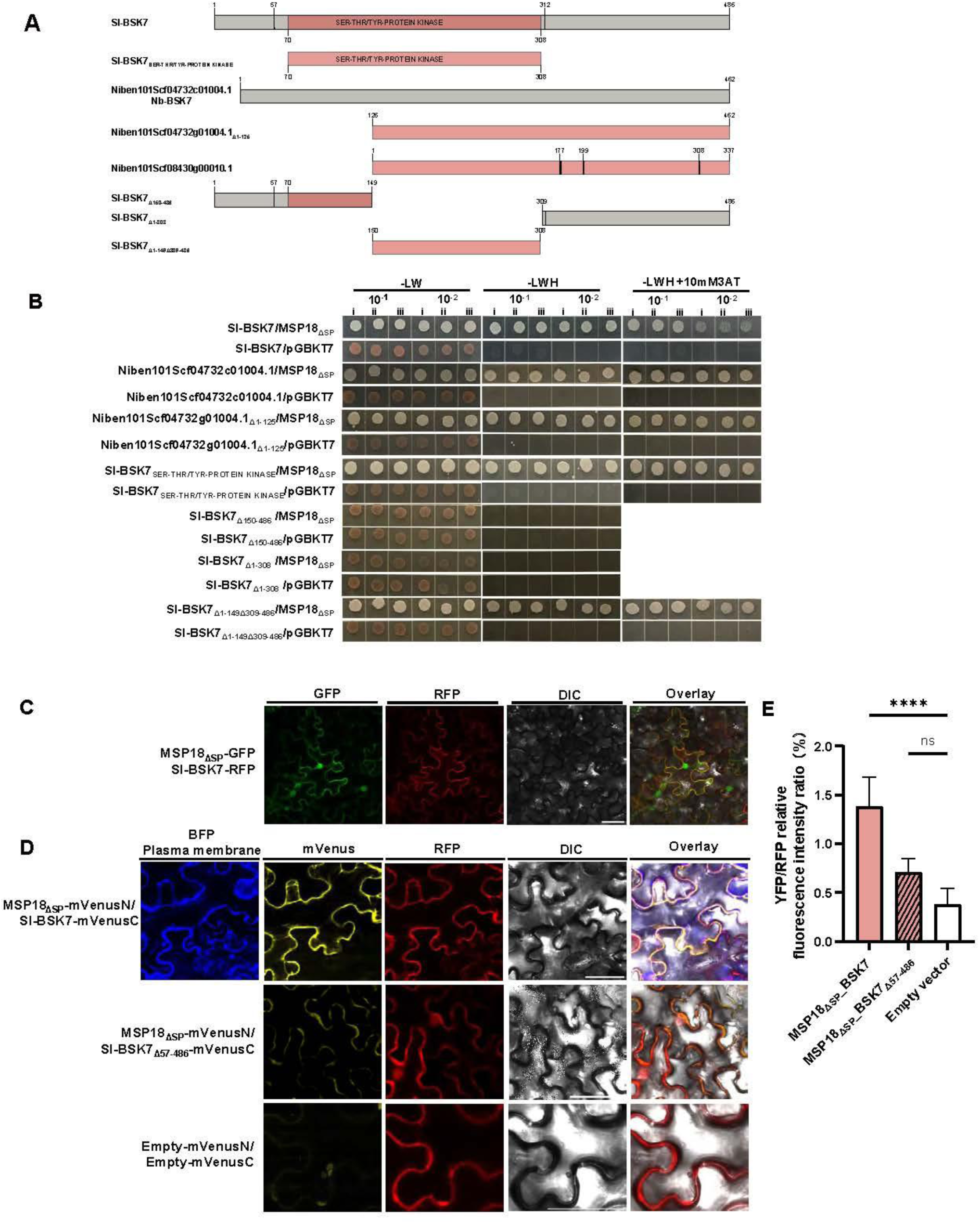
Mj-MSP18_ΔSP_ interacts with Sl-BSK7 in yeast and *in planta*. **(A)** Schematic representation of the domain organization and truncated versions of BSK7 homologs from different plant species. **(B)** Y2H analysis to refine the region of Sl-BSK7 required for interaction with Mj-MSP18_ΔSP_. PJ69-4A yeast cells were co-transformed with pGBKT7: Mj-MSP18_ΔSP_ and truncated versions of Sl-BSK7 or Nb-BSK7 (Niben101Scf04732c01004) in pGADT7. Transformants were spotted on SD medium lacking leucine and tryptophan (-LW) or additionally histidine (-LWH), with or without additional 3-AT. Autoactivation controls included co-transformations of each BSK7 construct with the empty pGBKT7 vector. Conserved amino acids of the two *N.benthamiana* orthologs are indicated by black lines. Dilution series are shown across the top; i, ii and iii represent independent biological replicates. **(C)** Colocalization assay of Sl-BSK7 and Mj-MSP18_ΔSP_ in *N. benthamiana* leaves. Sl-BSK7 and Mj-MSP18ΔSP were C-terminally fused to GFP and RFP, respectively, and co-expressed under the control of the constitutive CaMV35S promoter. Representative differential interference contrast (DIC) and fluorescence images of leaf tissues are shown. Scale bars = 50 µm. **(D)** Confocal images from a ratiometric BiFC (rBiFC) assay demonstrate interaction between Mj-MSP18_ΔSP_ and Sl-BSK7 at the plasma membrane. Split mVenus-tagged fusion proteins were co-expressed in *N. benthamiana* leaves. PIP2-BFP was used as a plasma membrane marker. No or weak signal was observed for the Mj-MSP18_ΔSP_-Sl-BSK7_Δ57-486_ combination. Negative controls with empty vector fusions showed only background and chlorophyll autofluorescence. RFP signal corresponds to a constitutively expressed control cassette. A total of 12–30 cells from four independent biological replicates were analyzed. Scale bars = 50 µm. **(E)** Quantification of the rBiFC (mVenus/RFP intensity ratio) reveals a significant difference between the Mj-MSP18_ΔSP_-Sl-BSK7 interaction and the negative control. The Mj-MSP18_ΔSP_–Sl-BSK7_Δ57–486_ combination did not yield a statistically significant increase over background. Data are presented as mean ± SEM from four biological replicates (n = 12–30; ****, *P* < 0.0001; one-way Brown-Forsythe and Welch ANOVA with Dunnett’s T3 multiple comparison test; ns = not significant).

### *In-planta* validation of the interaction between Mj-MSP18_ΔSP_ and Sl-BSK7

Following the identification of Sl-BSK7 as a proximal interactor by TurboID and the confirmation of direct interaction with Mj-MSP18_ΔSP_ in the Y2H assay, we next sought to validate this interaction *in planta*. A co-localization assay was performed in agroinfiltrated *N. benthamiana* leaf cells using fluorescently tagged versions of both proteins to examine whether they occupy similar subcellular compartments. As previously reported, Sl-BSK7 localizes to the plasma membrane (27). Consistent with this, we observed co-localization of Mj-MSP18_ΔSP_ with Sl-BSK7 at the plasma membrane, along with an additional nuclear localization of Mj-MSP18_ΔSP_ **(Figure 2C).**

To further confirm the interaction in planta and identify the subcellular site of interaction, a ratiometric bimolecular fluorescence complementation (BiFC) assay was performed (39). The construct included an internal red fluorescent protein (RFP) marker for expression control and allowed ratiometric comparison of fluorescence between experimental MSP18-BSK7 combinations and the negative controls. As a negative control, an N-terminal truncated version of Sl-BSK7 (Sl-BSK7_Δ57-486_), which retains the predicted myristoylation and palmitoylation sites but did not interact with Mj-MSP18 in the Y2H assay, was used along with Mj-MSP18_ΔSP_ in the same construct. Strong yellow fluorescence was detected from the MSP18_ΔSP_ -BSK7 BiFC pair, whereas no such signal was observed from the MSP18_ΔSP_ -BSK7_Δ57-486_ combination, despite similar RFP intensities. This indicates that Mj-MSP18_ΔSP_ specifically interacts with full-length Sl-BSK7 *in planta*. Co-localization of the BiFC signal with the plasma membrane marker (blue fluorescent protein, BFP) further confirmed that this interaction occurs at the plasma membrane (**Figure 2D**). Statistical analysis of the mVenus/RFP fluorescence intensity ratios revealed highly significant differences between the MSP18_ΔSP_/Sl-BSK7 pair and both negative controls—MSP18_ΔSP_/Sl-BSK7_Δ57–486_ and empty vectors—providing robust evidence for the specific interaction between the Mj-MSP18_ΔSP_ and Sl-BSK7 *in planta* (**Figure 2E**). These results collectively confirm a specific, plasma membrane-localized interaction between Mj-MSP18_ΔSP_ and full-length Sl-BSK7 *in planta*.

### Mj-MSP18 interferes with the BSK7-FLS2 interaction in yeast and *in planta*

Interestingly, BSK7 homologs from *A. thaliana* and *S. lycopersicum* have been shown to interact with the immune receptor FLS2 (25, 27). We found that the same truncated version of Sl-BSK7 that robustly binds MSP18_ΔSP_ was also both necessary and sufficient for a weaker interaction with the cytoplasmic domain of Sl-FLS2 (**Figure S7A, S7C**). As MSP18_ΔSP_ may competitively bind to and mask the shared binding region of Sl-BSK7, we hypothesized that its binding could interfere with the BSK7-FLS2 association.

To test this, we employed a yeast-3-hybrid (Y3H) assay, in which Mj-MSP18_ΔSP_ was introduced as a third protein to assess its effect on the Sl-BSK7 –Sl-FLS2 interaction. The Y3H results demonstrated that Mj-MSP18 _ΔSP_ disrupted the BSK7-FLS2 interaction in both tomato and *Arabidopsis* systems (**Figure 3A**). Residual yeast growth was occasionally observed, likely due to variation in effector expression levels. To mitigate this variability and quantitatively evaluate the disruption, the Y3H assay was performed on X-α-gal-containing medium (40). In the presence of Mj-MSP18_ΔSP_, reporter activity was consistently lost, indicating that BSK7–FLS2 interaction was effectively disrupted. A similar inhibitory effect was observed for the *Arabidopsis* AtBSK7-FLS2 and AtBSK8-FLS2 associations in when Mj-MSP18_ΔSP_ was co-expressed (**Figure 3B**). These results indicate that Mj-MSP18_ΔSP_ interferes with key components of PTI signaling complexes across plant species.

**Figure 3.**
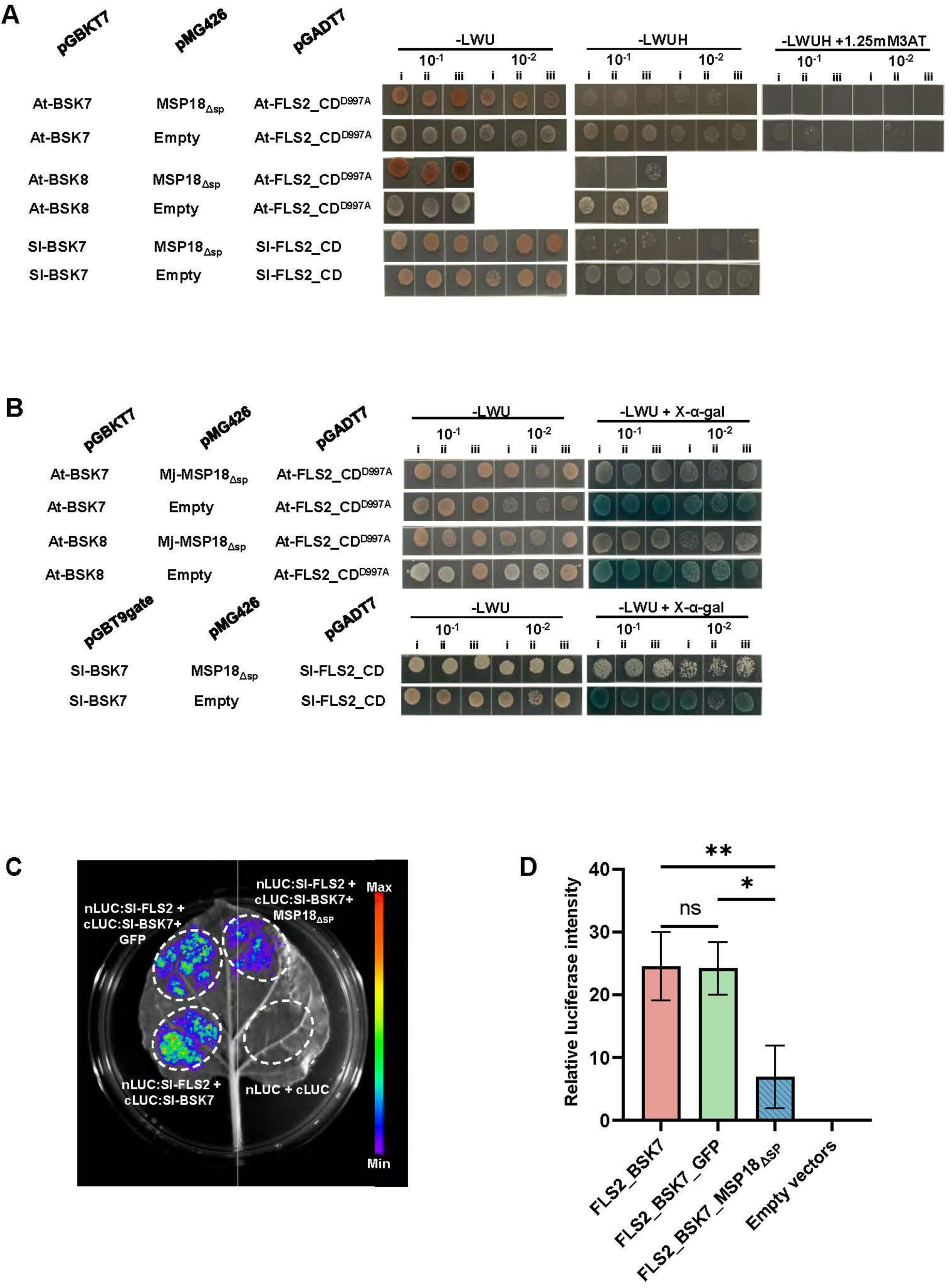
Mj-MSP18_ΔSP_ interferes with the BSK7-FLS2 interaction in yeast and *in planta*. **(A)** Y3H analysis with Sl-BSK7 and its orthologs from *A. thaliana* (At-BSK7 and 8) as baits, At-FLS2_CD (catalytically inactive form, At-FLS2_CD^D997A^; Majhi *et al.*, 2021) and Sl-FLS2_CD as preys, and Mj-MSP18_ΔSP_ as ‘bridge’ protein. Transformed yeasts were spotted in 10-fold and 100-fold dilutions on control medium (-Leu-Trp-Ura [-LWU]) and selective medium (-Leu-Trp-Ura-His [-LWUH]) with or without 3-amino-1,2,4-triazole (3-AT). Gene constructs in the pGBKT7 and pGADT7 vectors carry the GAL4-DNA-binding domain or the transcription activation domain, respectively, whereas pMG426 constructs do not contain any additional domain. **(B)** Y3H assay coupled with X-α-gal to quantitatively assess the role of Mj-MSP18_ΔSP_ in the BSK7-FLS2 interaction. The same constructs and combinations described in (A) were used, except that Sl-BSK7 was fused to the GAL4 DNA-binding domain in the pGBT9gate vector. Transformed yeast cells were spotted in indicated dilutions on control medium (-LWU) and chromogenic medium (-LWUH + X-α-gal) to detect interaction. For all yeast assays, 10x and 100x represent dilutions from overnight cultures. Labels i, ii, and iii represent three independent biological replicates. **(C)** *N. benthamiana* leaves were co-infiltrated with *Agrobacterium* containing constructs for expression of the indicated proteins fused to cLUC or cLUC, in the presence of Mj-MSP18_ΔSP_ or GFP. Sl-BSK7 and Sl-FLS2 served as a positive control, previously reported to interact in the luciferase complementation imaging assay (27). The empty vector combination (nLUC and cLUC) was used as a negative control. At 48 h post-infiltration, luciferase activity was visualized by imaging luminescence in whole leaves. The pseudocolor bar indicates the range of luminescence intensity. Bright and dark field images were overlaid to visualize the infiltrated leaf discs. Results represent four independent biological replicates. **(D)** Quantification of relative luciferase activity in leaves using Indigo imaging software. Each bar represents the mean ± SEM of four independent biological and three technical replicates. Asterisks indicate statistical significance as determined by one-way Brown-Forsythe and Welch ANOVA with Tukey’s post hoc test (*, P < 0.05; **, P < 0.01; ns, not significant).

A luciferase activity assay was conducted to validate the inhibition of the BSK7-FLS2 interaction by Mj-MSP18_ΔSP_ *in planta* (27). Co-expression of Mj-MSP18_ΔSP_ significantly reduced BSK7–FLS2–dependent luciferase activity (**Figure 3C**, **3D)**, confirming interference *in planta*.

Additionally, Y2H assays revealed potential homodimerization of At-BSK7, At-BSK8 as well as Sl-BSK7. A heterodimeric interaction between At-BSK7 and At-BSK8 was also observed **(Figure S7B)**.

### Mj-MSP18 compromises plant basal immunity by suppressing ROS production, callose deposition and HR

*Arabidopsis* BSK7 and BSK8, along with Sl-BSK7 interact with the FLS2 receptor and play important roles in immune signaling (25, 27). Given the ability of Mj-MSP18 to disrupt the BSK7-FLS2 association *in planta*, we sought to determine whether the effector could suppress flg22-triggered plant immune responses. To this end, ROS production induced by flg22 was measured in agro-infiltrated *N. benthamiana* leaves expressing Mp10 (41), eGFP, or Mj-MSP18_ΔSP_. At 72 hours post-infiltration, Mj-MSP18_ΔSP_ significantly reduced ROS generation compared to the GFP negative control (**Figure 4A**). Additionally, flg22-triggered callose deposition was analyzed in *N. benthamiana* leaves expressing Mj-MSP18_ΔSP_ or GFP. A significant decrease of callose deposits was observed in Mj-MSP18_ΔSP_-expressing tissues compared to GFP controls, further supporting its role in suppressing PTI responses. Notably, this effect was consistent across untagged Mj-MSP18ΔSP and both N- and C-terminal GFP fusions (**Figure 4B**).

**Figure 4.**
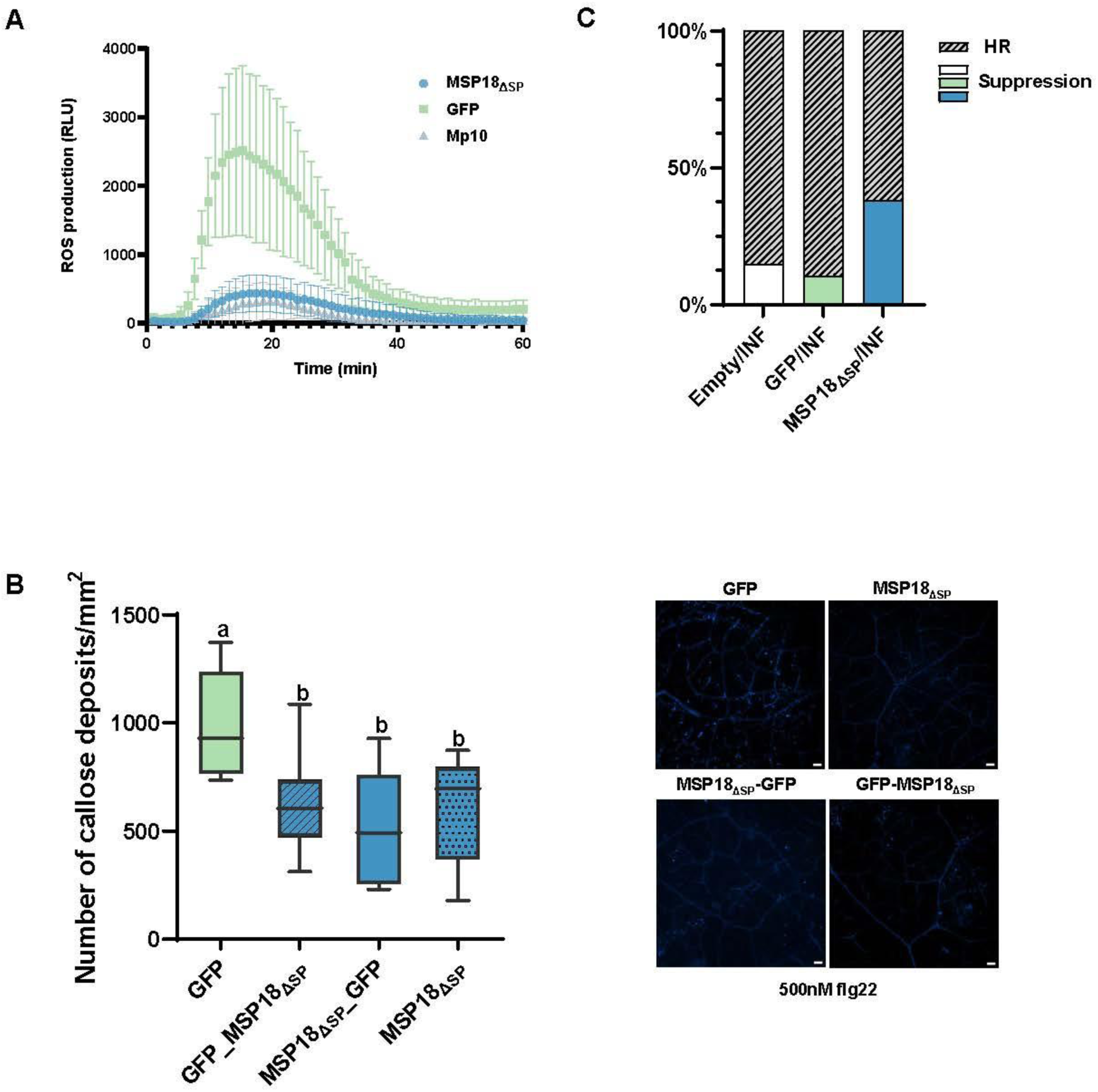
Mj-MSP18 suppresses plant basal immunity. **(A)** Mj-MSP18_ΔSP_ suppresses ROS accumulation in *N. benthamiana* leaves. Leaves transiently expressing Mj-MSP18_ΔSP_, GFP (negative control), or MP10 (positive control) were treated with 100 nM flg22 and analyzed for ROS production. **(B)** Callose deposition induced by 500 nM flg22 in *N. benthamiana* leaves transiently expressing Mj-MSP18_ΔSP_, N- and C-terminal GFP fusions of Mj-MSP18_ΔSP_, and GFP control. Representative images (bottom right) were captured at 24 h after flg22 treatment. Scale bar = 50 μm. Left: Quantification of callose spots per 1 mm^2^ area; means were calculated from 7-9 biological replicates. Different letters indicate significant differences between groups, as determined by one-way ANOVA followed by Tukey’s multiple comparison test at a 5% significance level. **(C)** Hypersensitive response (HR) suppression assays in *N. benthamiana*. *Mj-MSP18_ΔSP_* or GFP control was co-infiltrated with *INF1* in *N. benthamiana* leaves. The percentage of necrotic versus suppressed spots was scored at 48 h post-infiltration. A total of 60 infiltration sites were assessed per treatment. The experiment was repeated three times with similar results; one representative replicate is shown.

In addition, Mj-MSP18_ΔSP_ was also able to suppress localized cell death elicited by INF1 in *N. benthamiana* leaves (**Figure 4C**) (26), indicating broader suppression of defense-associated responses, including the hypersensitive response (HR).

### Effector-target protein BSK is involved in plant defense to root-knot nematode infection

As At-BSK7 and At-BSK8 in *A. thaliana* and Sl-BSK7 in *S. lycopersicum* were identified as targets of the RKN effector Mj-MSP18, we hypothesized that these proteins may influence nematode parasitism. To dissect the role of At-BSK7 and At-BSK8 in plant-nematode interactions, we assessed the susceptibility of *Arabidopsis* wild-type (WT) Col-0, *bsk7*, and *bsk8* single mutants, as well as *bsk7,8* double mutants, to *M. javanica*. Given that At-BSK7(8) is required for the BSK7(8)-FLS2 protein complex to function in plant basal immunity, *fls2* mutants were also included in the infection assay. Similarly, *Sl-bsk7* and *Sl-fls2* mutant tomato lines (42) were subjected to RKN infection alongside their corresponding wild-type backgrounds.

Our infection assays (**Figure 5A**, **5B**) demonstrated enhanced susceptibility in the *Arabidopsis fls2*, *bsk8* single mutants, and *bsk7,8* double mutants, as evidenced by a significantly higher number of galls observed at 10 days post-inoculation (dpi) and increased egg mass production at 6 weeks post-inoculation (wpi) compared to Col-0. The *bsk7* single mutant did not show a significant increase in gall or egg mass numbers in comparison to the control; however, the *bsk7,8* double mutant produced significantly more egg masses at 6 wpi than the *bsk8* single mutant.

**Figure 5.**
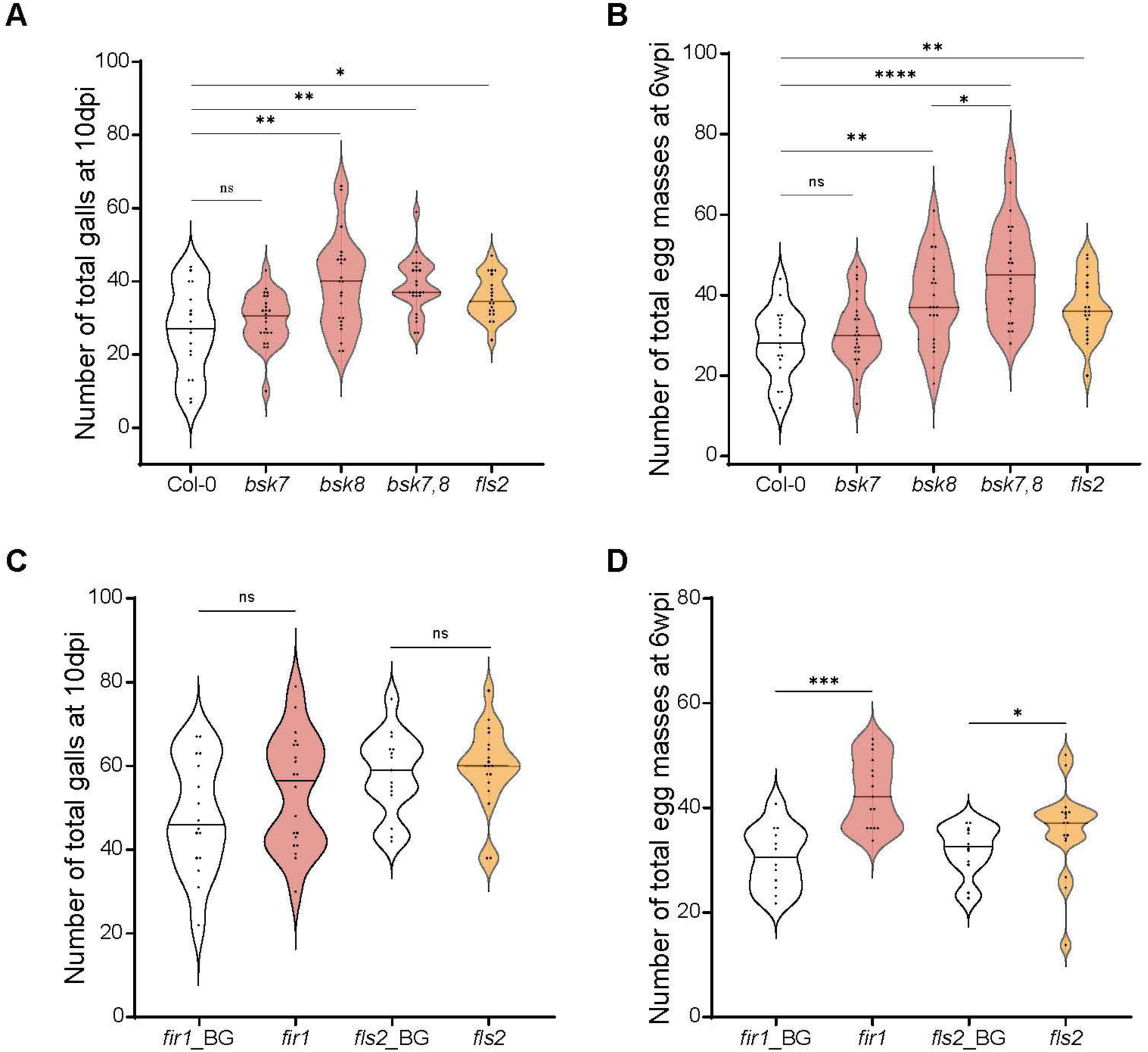
BSK7/8 orthologs are involved in plant defense to RKN infection. **(A)** Number of galls in *A. thaliana* at 10 days post inoculation (dpi) with *M. javanica*. The susceptibility of *Arabidopsis* wild-type Col-0, the *fls2*, *bsk7*, *bsk8* single mutants, and *bsk7,8* double mutants to *M. javanica* was compared. Inoculations were performed on 17-24 *in vitro* plants per treatment, each infected with 180 surface-sterilized J2s. Galls per plant were counted under a dissecting microscope at 10 dpi, and egg masses were counted at 6 weeks post-inoculation (wpi). Statistical analyses were performed using the Wilcoxon rank test in R to assess differences in susceptibility among genotypes; Prism was used for data visualization. Each dot represents an individual plant, and violin plots show data distribution. Statistical significance is denoted above comparisons between mutants and WT. Graphs represent data from three independent experiments. **(B)** Number of egg masses at 6 wpi. This panel presents the number of egg masses per plant produced by *M. javanica* on the *Arabidopsis* lines at 6 wpi. Statistical analyses and data representations are as described in (**A**). **(C)** Number of galls in *S. lycopersicum* at 10 dpi with *M. javanica*. The susceptibility of tomato *Sl-bsk7 (fir1)* and *Sl-fls2* mutants, along with their respective wild-type backgrounds (Hawaii 7981 and Rio Grande prf3), was assessed. Inoculations were performed on 15-20 *in vitro* plants per treatment, with each plant infected with 180 surface-sterilized J2s. Galls were counted at 10 dpi under a dissecting microscope, and egg masses at 6 wpi. Statistical analyses and data representation are as described in (**A**). **(D)** Number of egg masses at 6 wpi. This panel presents the egg mass counts on the same *S. lycopersicum* lines at 6 wpi. Statistical analysis and graphical presentation are as described in (**A**). Statistical significance is denoted above the comparisons, indicating differences between the mutants and the WT. Graphs represent data combined from three independent experiments. For all panels, non-significant differences are indicated as ns (*P* > 0.05), and significant differences are indicated with asterisks (*, *P* < 0.05; **, *P* < 0.01; ***, *P* < 0.001; ****, *P* < 0.0001).

The infection assay in tomato mutants did not show a significant difference in gall numbers at 10 dpi in *Sl-bsk7*(*fir1*) and *Sl-fls2* mutants compared to their corresponding wild-type lines, Hawaii 7981 and Rio Grande *prf3*, respectively (**Figure 5C**). Nevertheless, both mutants showed a significant increase in the egg mass production at 6 wpi compared to their respective WT controls (**Figure 5D**).

### Mj-MSP18 modulates the plant transcriptome

To broaden our comprehension of Mj-MSP18 function in plant cells, the impact on gene expression in tomato hairy root lines expressing Mj-MSP18_ΔSP_ or eGFP (**Figure S8A, S8B**) was examined. Analysis of differentially expressed genes (DEGs) 24 hours after estradiol induction identified 873 highly upregulated (*p*adj < 0.01; log₂ fold change > 1) and 1,415 highly down-regulated (*p*adj < 0.01; log₂ fold change < -1) genes in Mj-MSP18_ΔSP_-expressing lines compared to the eGFP control **(Supplemental Table 2)**. RT-qPCR validation of 10 selected genes confirmed the transcriptomic trends for 7 genes, including consistent up- or downregulation, while the remaining 3 (*CPD, BR6OX1*, and *MPK3*) showed similar but non-significant trends (**Figure S8C, Supplemental Table 3)**.

In Mj-MSP18_ΔSP_-expressing roots, gene ontology (GO) term enrichment analysis revealed significant downregulation of genes associated with “response to hydrogen peroxide” and “defense response” **(Figure 6A, Supplemental Table 4)** (26). In contrast, upregulated genes were significantly enriched for GO terms related to oxidoreductase activity, suggesting a shift in redox-related processes **(Figure 6B, Supplemental Table 5)**.

**Figure 6.**
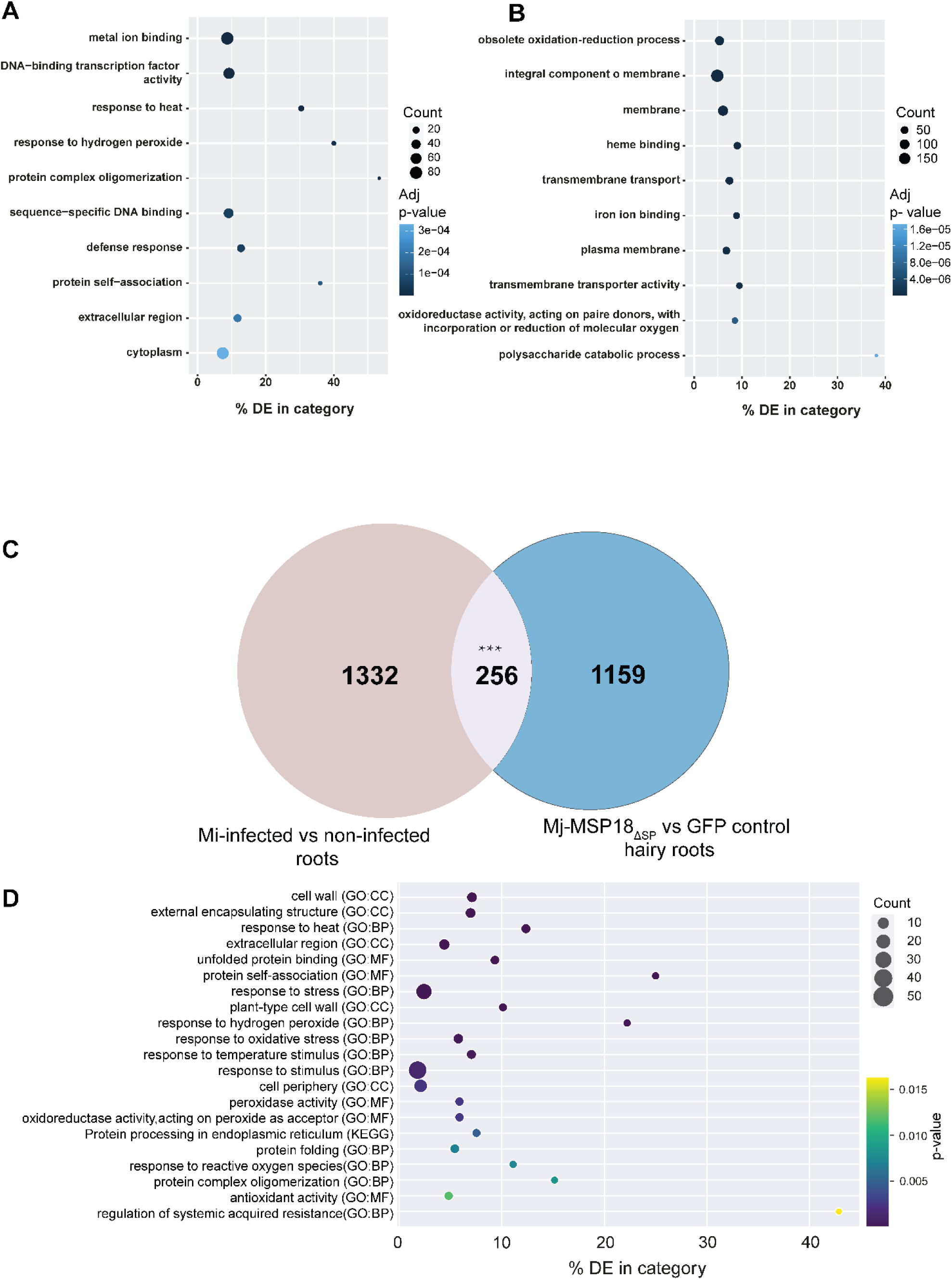
RNA-seq analysis reveals Mj-MSP18_ΔSP_ modulating the plant transcriptome. **(A)** Identification of differentially expressed genes (DEGs) by GO enrichment analysis in tomato hairy roots expressing Mj-MSP18_ΔSP_ compared to eGFP. RNA was extracted 24 hours after induction with 100 µM estradiol from four independently transformed hairy root lines. Bubble Plot shows the top ten overrepresented gene categories downregulated in Mj-MSP18_ΔSP_-expressing roots compared to eGFP (control; *P*-value < 0.01). **(B)** Bubble plot of top ten overrepresented gene categories significantly upregulated in Mj-MSP18_ΔSP_ compared to eGFP (control; *P*-value < 0.01). **(C)** Venn diagram comparing significant DEGs between *M. incognita-*infected vs uninfected tomato roots and Mj-MSP18_ΔSP_ vs GFP-overexpressing tomato hairy roots. (*P*-value < 0.01). Asterisks indicate a significant overlap (*** *P* < 0.001; Chi-squared test performed in R, X-squared = 720.79, df = 1, *P*-value < 2.2e-16). (D) Bubble plot showing the top twenty overrepresented GO categories among the 256 commonly significantly downregulated genes in both *M. incognita* infected vs uninfected tomato roots (111) and Mj-MSP18_ΔSP_ vs GFP-expressing tomato hairy roots. GO enrichment was performed using the g:Profiler (g: Goast) tool; plotting was done using Python (3.9) and Matplotlib plotting library.

To gain deeper insights into shared biological functionality between Mj-MSP18-expressing hairy roots and naturally RKN-infected tomato roots, a comparative transcriptome analysis was performed. Differentially expressed genes from Mj-MSP18 vs. eGFP hairy roots were compared with publicly available gene expression profiles of *M. incognita* infected versus uninfected tomato roots. This revealed commonly down-regulated genes enriched in GO categories such as “response to stress”, “peroxidase activity”, “response to reactive oxygen species”, and “regulation of systemic acquired resistance” among others **(Figure 6C**, **6D and Supplemental Table 6)**.

### Y2H screen for additional nuclear plant targets of Mj-MSP18

As aforementioned, several significantly enriched candidate proteins were identified in Mj-MSP18 samples using TurboID **(Supplemental Table 1).** One of the top hits, Sl-BSK7, was confirmed as an Mj-MSP18 _ΔSP_-interacting partner localized to the plasma membrane. Given that subcellular localization assays revealed an additional nuclear localization of Mj-MSP18_ΔSP_ **(Figure S2)**, and since no nuclear proximal partners were identified using TurboID, we sought to specifically identify potential nuclear plant targets of the effector.

To this end, we performed a Y2H screen (43), using Mj-MSP18_ΔSP_ fused to the GAL4-DNA-binding domain as bait, and a tomato cDNA library (44) fused to the GAL4 activation domain as prey (45). The Y2H screen yielded 10 candidate interactors **(Supplemental Table 7);** however, only MYB (*Solyc03g113620*) and MYC2 (*Solyc08g076930*) could be validated in follow-up pairwise Y2H assays using full-length coding sequences. These two transcription factors consistently interacted with Mj-MSP18_ΔSP_ in yeast **(Figure 7A)**, whereas other candidates failed to validate, likely due to artifacts such as truncated clones, non-specific interactions.

**Figure 7.**
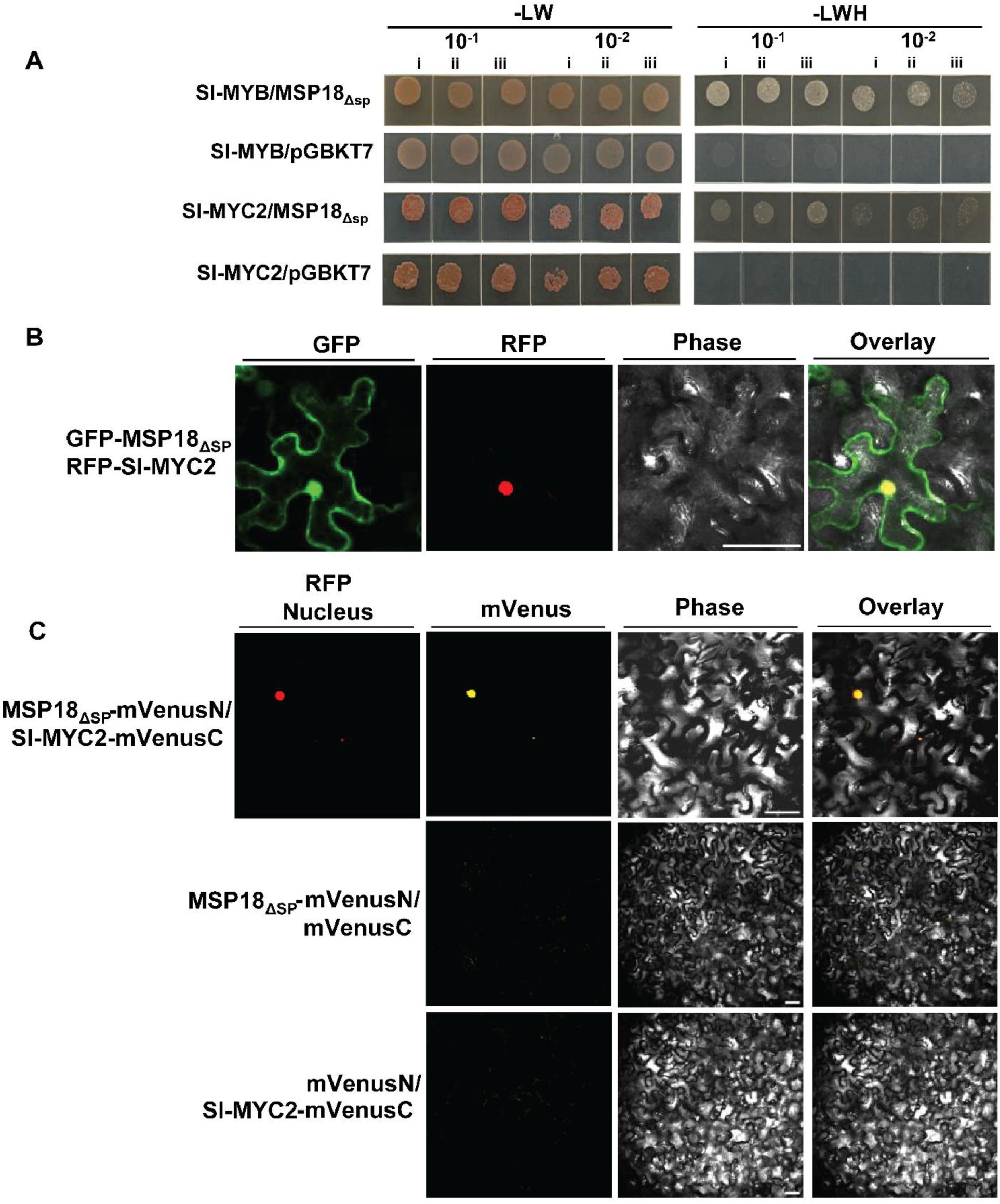
Interaction between Mj-MSP18_ΔSP_ and two plant transcription factors found by Y2H. **(A)** Detection of interaction between Mj-MSP18_ΔSP_ and Sl-MYB (Solyc03g113620) or Sl-MYC2 (Solyc08g076930) by Y2H assay. PJ69-4A yeast cells were co-transformed with pGBKT7::Mj-MSP18_ΔSP_ and either pGADT7::Sl-MYB or pGADT7::Sl-MYC2. Cells were spotted on SD (Synthetic Defined) medium lacking leucine and tryptophan (-LW) or leucine, tryptophan, and histidine (-LWH), with or without additional 3-amino-1,2,4-triazole (3-AT). Autoactivation controls included yeast transformed with pGADT7::Sl-MYB or pGADT::Sl-MYC2 and theempty pGBKT7 vector. Dilution series are indicated at the top; i, ii, and iii represent independent biological replicates. **(B)** Mj-MSP18_ΔSP_ and Sl-MYC2 show co-localization in plant cells. Representative differential interference contrast (DIC) and fluorescence images of *N. benthamiana* leaves transiently expressing *Mj-MSP18_ΔSP_-eGFP* or *Sl-MYC2*. Scale bars = 50 µm. **(C)** Nuclear interaction between Mj-MSP18_ΔSP_/Sl-MYC2 detected by bimolecular fluorescence complementation (BiFC) in *N. benthamiana* leaves transiently expressing the indicated constructs along with a nuclear localization marker. Arabidopsis fibrillarin protein FBR1 tagged with RFP was used as a nuclear marker. Negative BiFC controls are shown in the middle and lower panels. Scale bars = 50 µm.

To validate these interactions in planta, co-localization and BiFC assays were performed in *N. benthamiana* leaves **(Figure 7B**, **7C)**. Subcellular localization showed that Mj-MSP18_ΔSP_ and MYC2, tagged with eGFP and RFP respectively, co-localize in the nucleus (46). In the BiFC assay, co-expression of mVenusN –Mj-MSP18_ΔSP_ and mVenusC –Sl-MYC2 resulted in reconstituted fluorescence, suggesting *in planta* interaction nuclear co-localization, as further confirmed by signal overlap with the nuclear marker FIBRILLARIN 1 –RFP mrker **(Figure 7C)**.

Attempts to detect RFP fluorescence from the Sl-MYB –RFP fusion construct were unsuccessful (data not shown). Although this may indicate low expression or instability of the MYB fusion protein in *N. benthamiana*, further experiments are required to determine the cause.

### Differentially expressed genes (DEGs) induced by Mj-MSP18 harbor MYB/MYC2 binding motifs in their promoters

Given that Mj-MSP18_ΔSP_ interacts with the MYB and MYC2 transcription factors, we hypothesized that it may influence their downstream transcriptional targets *in planta*. Therefore, we re-analyzed the DEGs induced by Mj-MSP18_ΔSP_ compared to GFP expression in tomato hairy root lines and examined their promoter regions (3 kb upstream of the start codon) for potential MYB- and MYC2-binding sites (5’-AGATAA-3’ and 5’-CACGTG-3’, respectively) (47, 48). Enrichment analysis revealed that MYB- and MYC2-binding sites were significantly overrepresented in the promoters of DEGs induced by Mj-MSP18_ΔSP_ expression compared to their average occurrence in the tomato genome. Specifically, this background average refers to the proportion of all genes in the genome that harbor these binding sites. Notably, 81% of the highly upregulated DEGs harbored at least one MYB-binding site in their promoter region, and 32% harbored a MYC2-binding site **(Figure 8A**, **8C)**. GO term analysis of the significant DEGs containing MYB or MYC2 binding motifs **(Figure 8B**, **8D and Supplemental Table 8, 9)** revealed an overrepresentation of genes associated with the GO categories “cellular response to hormone stimulus” and “hormone-mediated signaling pathway”, which sparked our further exploration of the impact of Mj-MSP18_ΔSP_ expression on phytohormone levels *in planta*.

**Figure 8.**
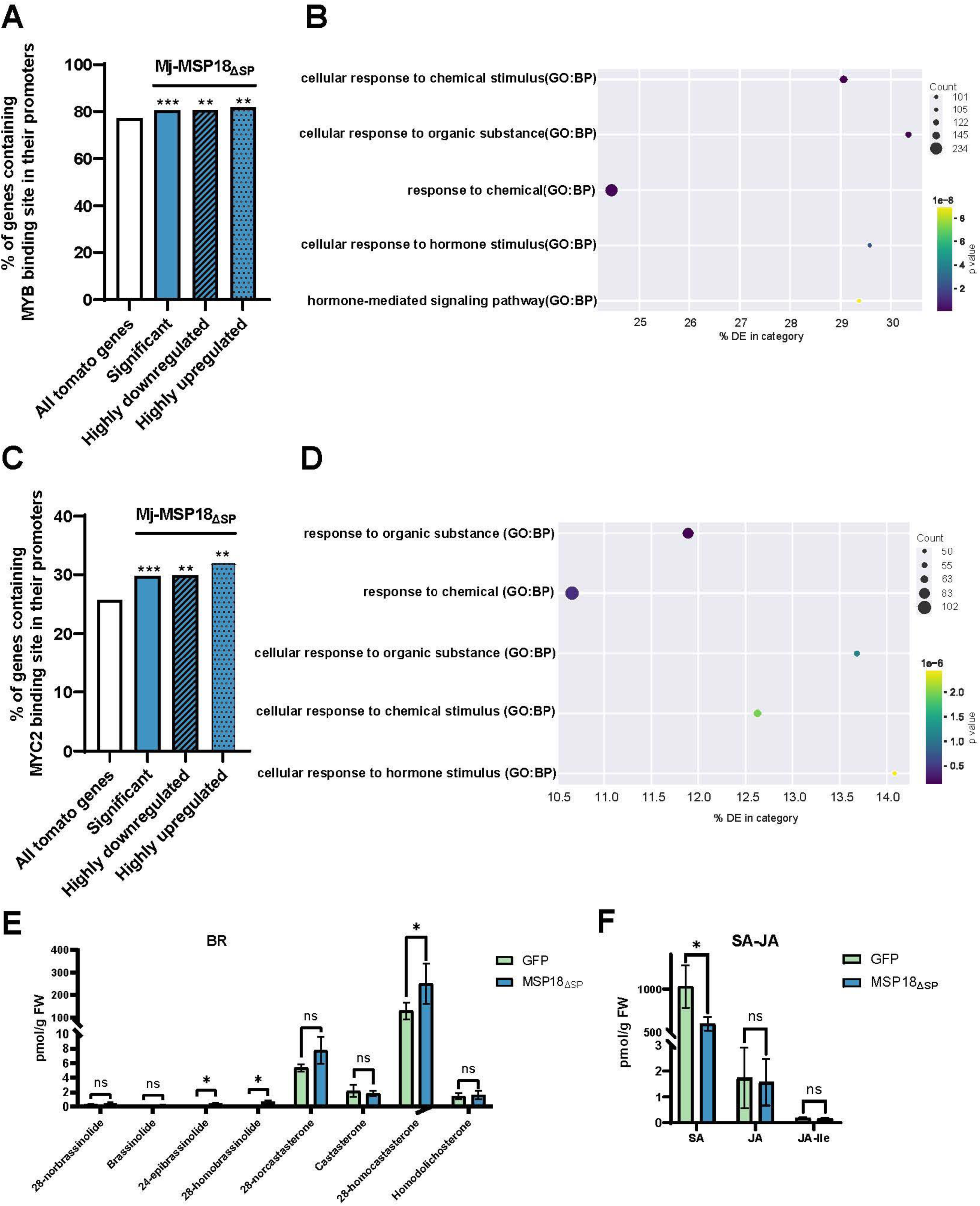
Motif analysis of Sl-MYB/MYC2 transcription factor binding sites in the Mj-MSP18_ΔSP_-induced transcriptome and corresponding metabolite validation. **(A)** Frequency of the MYB binding motif (5’-AGATAA-3’) in the promoter regions (3 kb upstream of the start codon) of Mj-MSP18_ΔSP_ differently expressed genes compared to all tomato genes. Differently expressed genes are categorized as: significantly different (*p*adj < 0.05), highly upregulated (log2FC >1 and *p*adj < 0.01) or highly downregulated (log2FC <-1 and *p*adj <0.01). Asterisks indicate significant enrichment of the binding site (** *P* < 0.01, *** *P*< 0.001; Fisher’s exact test). Analyses and plotting were performed using Python (3.9) with Biopython and Matplotlib. **(B)** Bubble plot showing the top five overrepresented gene categories of 4,170 significant DEGs containing the Sl-MYB binding site in Mj-MSP18_ΔSP_ vs GFP (5% significance level; see **Supplemental Table 8)**. GO enrichment was performed using the g:Profiler (g:GOSt) tool; visualization used Python (3.9) and Matplotlib. Abbreviations: GO, Gene Ontology; KEGG, Kyoto Encyclopedia of Genes and Genomes; BP, Biological Process; DE, differentially expressed. **(C)** Frequency of MYC2 binding motif (5’-CACGTG-3’) in promotr regions (3 kb upstream of the start codon) of Mj-MSP18_ΔSP_ differentially expressed genes compared to all tomato genes. Categories and statistical tests are as described in (**A**). **(D)** Bubble Plot showing the top five overrepresented gene categories among 1,546 significant DEGs containing the MYC2 binding site in Mj-MSP18_ΔSP_ vs GFP (5% significance level; see **Supplemental Table 9)**. g: profiler (g: Goast) tool was used for analysis; Python (3.9) and Matplotlib plotting library were used for plotting. GO enrichment and plotting tools were described in (**A**). **(E)** and **(F)** Quantification of phytohormone levels (BRs, SA, JA and JA-Ile) in *Mj-MSP18_ΔSP_* and *GFP* expressing tomato hairy roots following 24-hour estradiol induction. Data are presented as means ± standard deviation (SD) from four biological repeats (n = 4). Statistical significance was determined using an unpaired two-sample Student’s *t*-test. Non-significant differences are indicated as ns (*P* > 0.05); significant differences are marked with an asterisk (*, *P* < 0.05). JA, jasmonic acid; JA-Ile, (-)-jasmonyl-L-isoleucine; SA, salicylic acid; BRs, brassinosteroids.

**Figure 9.**
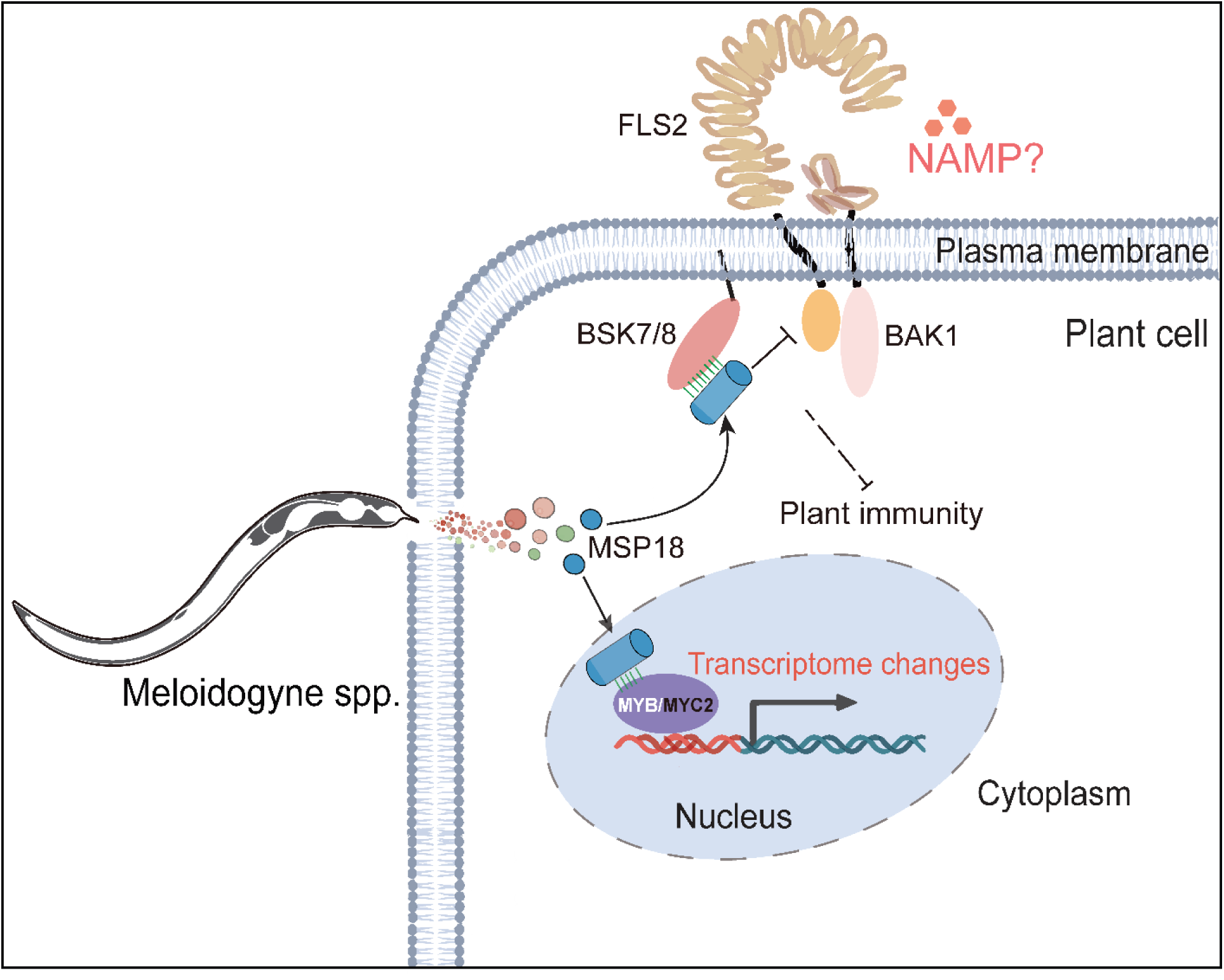
Model for the function of Mj-MSP18 during plant–nematode interaction. Root-knot nematodes (RKNs) secrete the effector Mj-MSP18_ΔSP_ from their pharyngeal glands into plant cells, where it interacts with BSK7/8 at the plasma membrane. The interaction interferes with the association between BSK7/8 and FLS2, thereby suppressing flg22-induced pattern-triggered immunity (PTI) responses, including reactive oxygen species (ROS) production and callose deposition. Additionally, Y2H identified interaction between Mj-MSP18_ΔSP_ and the transcription factors MYB and MYC2 in the nucleus, potentially contributing to the transcriptomic changes observed in tomato hairy roots.

### Targeted metabolite analysis reveals the involvement of Mj-MSP18 in the regulation of hormone biosynthesis

Many MYB-regulated genes associated with the GO categories “brassinosteroid biosynthetic process” and “jasmonic acid biosynthetic process” were significantly upregulated. In contrast, genes involved in SA biosynthesis regulation were significantly downregulated **(Supplemental Table 8)**. A similar pattern was observed among MYC2-targeted differentially expressed genes (DEGs) in the Mj-MSP18_ΔSP_-expressing samples **(Supplemental Table 9)**.

A targeted metabolite analysis was performed to quantify jasmonic acid (JA), salicylic acid (SA), and brassinosteroid (BR) levels in Mj-MSP18_ΔSP_-3xHA and eGFP-3xHA tomato hairy root samples following 24 hours of estradiol induction. Mj-MSP18_ΔSP_ expression led to a significant increase in brassinosteroid levels, including 28-norcastasterone and 28-homocastasterone, and a reduction in SA levels, while JA levels remained unchanged **(Figure 8E**, **8F).**

## Discussion

Many biotrophic pathogens challenge plants, and nematodes are no exception. Like other pathogens, root-knot nematodes deploy an arsenal of effector proteins to subvert plant defenses (49, 50). In our study, we focused on the RKN effector Mj-MSP18 and uncovered its role in manipulating plant immunity. Our results demonstrate that Mj-MSP18, which is highly conserved among *Meloidogyne* species, functions as a molecular “Swiss army knife” by targeting the plant kinase BSK7 at the plasma membrane as well as MYC and MYB transcription factors. Through these interactions, Mj-MSP18 reprograms the cell’s metabolic and transcriptomic state, leading to multifaceted immunity suppression. We identified and confirmed these interactions using TurboID proximity labeling, yeast two-hybrid (Y2H), and *in planta* validation assays, which showed that Mj-MSP18 disrupts the formation of the BSK7–FLS2 complex—a critical hub in pattern-triggered immunity (PTI) known for its role in bacterial and fungal defense (25). Although nematodes do not produce flagellin, it is tempting to speculate— based on our infection results and recent evidence for the discovery of new NAMPs (51) that plants have receptors, including FLS2 or related proteins, to detect nematode-associated molecular patterns (NAMPs) (12, 13). Thus, Mj-MSP18’s interference with the BSK7–FLS2 complex might compromise both bacterial and nematode recognition, offering nematodes a sophisticated strategy to overcome early immune responses. Supporting this, the ability of Mj-MSP18 to suppress ROS burst and defense aligns with prior infection assays performed on plants overexpressing its homolog Mi-MSP18, which similarly enhanced parasitism while suppressing defense responses (26).

Our transcriptomic analysis of tomato hairy roots expressing Mj-MSP18_ΔSP_ revealed extensive reprogramming of the host gene expression profile. Genes involved in oxidative stress responses and defense were significantly downregulated, while those associated with oxidoreductase activity and BR biosynthesis were upregulated. In particular, the upregulation of brassinosteroid (BR) biosynthetic genes **(Supplemental Table 8**), together with elevated BR levels in our metabolite analyses, suggests that Mj-MSP18 may shift the growth defense balance in favor- of nematode parasitism (26, 52, 53). The role of BRs in plant defense is unclear and might depend on the specific plant-pathogen interaction, the timing, the concentration and the type of BR.

Several studies demonstrated that BRs suppress PTI and enhance susceptibility (54, 55). By antagonism with JA-activated defense, BR enhanced susceptibility in rice to RKN (52) and the brown planthopper (56) and in Arabidopsis to beet armyworms (*Spodoptera exigua*) and the fungus *Botrytis cinerea* (57). On the other hand, BR induced defense to RKN in tomato (58) and to cyst nematodes in soybean (59) and potato (60).

The downregulation of salicylic acid (SA) biosynthesis—SA that confers resistance to root-knot nematodes (61) —further supports the idea that Mj-MSP18 reprograms hormone-mediated defense networks (5). Interestingly, in *Pythium graminicola*, a pathogen of rice, BRs were shown to suppress SA-mediated defenses via a decoy strategy acting downstream of SA biosynthesis (62), raising the question whether a similar mechanism is exploited by RKNs through Mj-MSP18-mediated BR signaling modulation. For example, receptor NILR1 has been reported to mediate both PTI and BR responses (63), which may underlie why nematode-associated molecules such as *Ascr18* inhibit root growth in potato but not in Arabidopsis (63). This species-specific dual role of NILR1 in potato, but not Arabidopsis, could explain differing susceptibilities and highlights that BR–PTI crosstalk during nematode infection may vary across plant species. Whether Mj-MSP18 interference with BSK signaling differentially affects these intertwined pathways depending on host species remains an intriguing hypothesis. Although no significant difference in jasmonic acid accumulation was observed between Mj-MSP18_ΔSP_ and GFP hairy root samples at 24 hours post-induction, additional time points could be analyzed (64, 65).

While multifunctional effectors like RHA1B (63, 66), have been described that target both membrane-localized receptors such as NILR1 and nuclear-regulated transcripts like *CycA2* mRNA, Mj-MSP18 is, to our knowledge, the first root-knot nematode (RKN) effector shown to directly interact with both plasma membrane-localised kinases and nuclear transcription factors, thereby coordinating hormone- and immunity-related gene expression. In our study, we found that Mj-MSP18 interacts with BSK7 at the plasma membrane and with nuclear transcription factors MYB and MYC2. These interactions, confirmed by Y2H and BiFC assays, suggest that Mj-MSP18 reprograms the plant cell’s metabolic and transcriptomic state at multiple levels, leading to a broad immune suppression. Importantly, we observed that the effector does not alter BSK7 localization, indicating that its mode of action involves disrupting the formation and function of signaling complexes rather than protein relocalization.

To further explore the relevance of the nuclear interactions, we examined whether the effector influences gene expression through MYB and MYC2. While this enrichment suggests potential direct regulation, we caution that motif presence alone does not confirm direct targeting. An alternative hypothesis is that the transcriptome reprogramming could be to some extent a downstream consequence of the disrupted BSK–FLS2 signaling complex, whereby Mj-MSP18’s interference with early membrane-associated immune signaling cascades indirectly rewires nuclear transcription. Therefore, both direct transcription factor modulation and indirect signaling disruption may contribute in parallel to the observed changes. Functional validation using ChIP-seq or promoter-reporter assays will be essential to confirm direct regulatory links. Together, these findings support a dual-compartment mode of action for Mj-MSP18 namely disruption of membrane-localized immune complexes while modulating nuclear transcription to exert comprehensive control over host defense.

Despite these advances and our understanding of which domain is targeted by Mj-MSP18 in different BSKs, several critical questions remain. First, the precise molecular mechanism by which Mj-MSP18 disrupts the BSK7–FLS2 interaction is still unclear. Is the disruption due to steric hindrance, an allosteric change in BSK7’s conformation, or might Mj-MSP18 recruit additional factors that destabilize the complex? High-resolution structural studies, such as cryo-electron microscopy could yield significant advances. In addition, assessing the phosphorylation status of both BSK7 and FLS2, as well as downstream components of MAPK signaling, is essential for understanding how signal transduction is affected by this disruption. Second, while our transcriptomic and hormonal data indicate a rebalancing of BR and SA pathways, the temporal dynamics of these changes and their causal relationships with immune suppression during RKN infection warrant further investigation. Longitudinal studies that capture hormone fluctuations and transcriptional responses over time would help elucidate these dynamics.

Moreover, our investigation of plant mutants corroborates the role of the BSK7–FLS2 axis in defense. Our results show that both *bsk7/8* and *fls2* mutants exhibit increased susceptibility not only to bacteria and fungi but also to nematodes. This observation supports the notion that FLS2’s role extends beyond flagellin recognition and may include the perception of nematode-associated molecular patterns (NAMPs). *bsk7* mutant lines also underscored its pivotal role in defense against root-knot nematodes. However, it remains to be determined whether the enhanced susceptibility in these mutants is solely due to disruption of BSK7–FLS2 signaling or if other indirect effects contribute. In addition, the identification of ESCRT complex components (e.g., AT5G16880 and AT1G21380) in our TurboID experiments suggests that endocytosis of FLS2 might also modulate its activity—a mechanism proposed to affect receptor recycling and signaling (67, 68). Whether these components influence the efficacy of the BSK7–FLS2 complex and, consequently, the plant’s overall immune status is an exciting avenue for future research. Time course experiments using fluorescent fusions of multiple proteins will be essential to confirm this potential internalization and to determine whether flg22 is required to downregulate this signaling and trigger internalization.

In addition to these molecular interactions, the long-term physiological impacts of disrupting the BSK7– FLS2 axis merit further investigation. Given the central role of these proteins in balancing growth and defense (18, 21), chronic interference by Mj-MSP18 and pleiotropic effects on plant fitness, resource allocation, and even the composition of the plant microbiome should be tested further.

On the other hand, BSK family members have also been shown to interact with NILR1—the LRR-RLK responsible for recognizing nematode-derived ascarosides—raising the possibility that the observed targeting of BSK7/8 by Mj-MSP18 might interfere not only with FLS2-BSK complexes but also with NILR1-BSK signaling (69). This suggests an alternative hypothesis: MSP18 may primarily disrupt NILR1–BSK interactions rather than FLS2–BSK complexes, a possibility that warrants further investigation.

In conclusion, our study demonstrates that the nematode effector Mj-MSP18 subverts plant immunity by targeting the BSK7–FLS2 complex, thereby dampening early PTI responses and reprogramming both the transcriptome and hormone signaling networks. This multifaceted mechanism, which also involves interactions with nuclear transcription factors such as MYB and MYC2, underscores the sophisticated strategies evolved by nematodes to overcome host defenses. Moreover, the possibility that FLS2 may recognize nematode-associated molecular patterns expands the functional repertoire of this receptor and indicates a potentially missing part of crosstalk between BR signaling and PTI. Addressing these unresolved questions through advanced structural studies, longitudinal analyses of hormone dynamics, and functional dissection of mutant phenotypes will pave the way for innovative strategies to bolster crop resistance to nematode infections, ultimately enhancing agricultural sustainability.

## Materials and Methods

### Strains

*M. javanica* and *M. incognita* morelos were maintained in cultures using susceptible tomato plants (*S. lycopersicum* cv Moneymaker) in two-liter plastic pots filled with potting soil (Structural, Kaprijke, Belgium). The infected tomato plants were grown in a greenhouse under 8/16-hour night/day conditions at 22–24 °C and 16 °C, respectively. *N. benthamiana Domin.* plants were grown in a greenhouse under 8/16-hour night/day conditions at 28 °C and 16 °C, respectively. *A. thaliana Heynh* plants were grown in a greenhouse under 8/16-hour night/day conditions at 20–22 °C and 16 °C, respectively. *Escherichia coli* Top10 and *Agrobacterium tumefaciens* GV3101 or *A. rhizogenes* ATCC 15834 were routinely grown in LB/YEB medium at 37 °C and 28 °C, respectively. The yeast strain *Saccharomyces cerevisiae* PJ69-4α was cultured at 30 °C for Y2H assays as described below. All plant extracts, tissues, bacterial and yeast strains were stored at -80 °C.

### Plasmid and strain construction

All plasmids and constructs used in this study are listed in **Supplemental Table 10**. Standard molecular cloning methods were used, and DNA assembly was performed using Gateway cloning, Gibson assembly or site directed mutagenesis. All oligos used in this study are listed in **Supplemental Table 11**. All constructed plasmids were checked for correctness by Sanger sequencing. Tomato *Sl-BSK7* (Solyc12g099830), tobacco *Nb-BSK7* (Niben101Scf04732c01004), *Arabidopsis At-BSK7* (AT1G63500) and *At-BSK8* (AT5G41260) coding sequences were used as templates for amplification from cDNAs.

### *In silico* sequence analysis

Using Meloidogyne INRAE Blast (https://meloidogyne.inrae.fr/), we found the closest homologue of *Mi-MSP18* (NCBI Accession number AAN08584) (32) to be Mjav1s11723g062659 (*M. Javanica* scaff11723g062659), here referred to as *Mj-MSP18*. Local alignment was done according to Smith– Waterman algorithm, and global alignment using Needleman–Wunsch algorithm, both implemented in Snapgene 6.0. SignalP 5.0 (70) was used to predict signal peptides. Mj-MSP18 without signal peptide is referred to as Mj-MSP18_ΔSP_. For all experiments, LOCALIZER (71) was used to predict protein localization. Motif scan (https://myhits.sib.swiss/cgi-bin/motif_scan), NCBI Conserved Domain Search (https://www.ncbi.nlm.nih.gov/cdd/) (72), and InterProscan (https://www.ebi.ac.uk/interpro/) (73) were used to predict motifs and domains in the proteins.

### Immunoblotting for TurboID

Hairy root cultures were harvested, flash-frozen in liquid nitrogen and stored at -80 °C until further processing. Tissue samples were ground in liquid nitrogen using a mortar and pestle, and the powdered material was transferred to a 15 or 50 mL Falcon tube. For each sample, 2.5 g of ground material (equivalent to volume of ∼500 µL) was resuspended in2 mL of urea-based high stringency homogenization buffer (100 mM Tris pH 7.5; 2% SDS; 8M urea; 150 mM NaCl), resulting in∼2.5 mL of lysate. Proteins were extracted by three cycles of freeze-thawing in liquid nitrogen. Lysates were cleared by centrifugation at 16,000 × *g* for 15 minutes, and the supernatants were transferred to fresh 15 mL tubes. This centrifugation step was repeated twice. To remove excess biotin, a desalting step was performed using Sephadex G-25 in PD-10 Desalting Columns (GE healthcare lifesciences, cat n° 17085101). For each setup, 400 μL of streptavidin-agarose bead slurry (50% v/v; Novagen, cat. nr 69203) was pre-washed three times with 2 mL of high-stringency BioID lysis buffer by centrifugation (600 × *g*, 5 min), followed by supernatant removal. To 4 mg of protein lysate, pre-washed beads were added, and biotinylated proteins were captured by overnight incubation with overhead rotation at room temperature. After incubation, the beads were pelleted (600 × *g*, 2 min), and the unbound supernatant was removed and retained for Western blotting. Beads were transferred to low-protein-binding tube (Eppendorf) and washed extensively: four times with 1 mL of high-stringency buffer (5 min each), followed by a 30 min wash with high-stringency buffer, a 30 min wash with high-salt buffer (1 M NaCl, 100 mM Tris-HCl, pH 7.5), and a final 5 min wash with 1 mL ultrapure water. Following the final wash, beads were divided into two parts: 90% for downstream processing and 10% for Western blot analysis. To 70 µL each of the input, desalted input, unbound, and bound fractions, 25 µL of sample loading buffer and 5 µL of reducing agent (both Bio-Rad) were added. Samples were then separated by SDS-PAGE (1.0 mm thick 4 to 12% polyacrylamide Criterion Bis-Tris XT-gels, Bio-Rad) in MOPS buffer (Bio-Rad) at 150 V. Subsequently, proteins were transferred onto a PVDF membrane. Membranes were blocked for 30 min in a 1:1 mixture of Tris-buffered saline (TBS) and Odyssey blocking solution (cat no. 927-40003, LI-COR, Lincoln, NE, USA) and probed by Western blotting. Following overnight incubation with primary antibodies in TBS-T (0.1% Tween-20)/Odyssey Blocking buffer and three 10-minute washes in TBS-T, membranes were incubated with secondary antibody for 30 minutes in the same buffer followed by 3 washes in TBS-T or TBS (last wash step). The following detection reagents and antibodies were used: Streptavidin, Alexa Fluor™ 680 Conjugate (Invitrogen, S32358, 1:10,000), mouse anti-FLAG (Sigma, F3165; 1:2,000), and anti-mouse (IRDye 800 CW goat anti-mouse antibody IgG, LI-COR, cat no. 926-32210, 1:10,000. Bands were visualized using the Odyssey Infrared Imaging System (LI-COR).

### Proteomics sample preparation after TurboID-mediated proximity labeling

As described above, following the final wash, the beads were split 90%/10%, and the supernatant was removed. Then, 90% of the beads were washed 3 × 5 min with 1 mL 50 mM ammonium bicarbonate (pH 8.0), resuspended in 250 µL of 50 mM ammonium bicarbonate (pH 8.0), and 1 μg of mass spectrometry grade trypsin (Promega, Madison, WI), dissolved in 250 μL of 50 mM ammonium bicarbonate (pH 8.0), was added. The samples were incubated overnight at 37 °C with vigorous mixing (850 rpm) to keep the beads in suspension. After overnight digestion, an additional 0.5 μg of trypsin was added, followed by a further2-hour incubation. The beads were pelleted (600 × *g*, 2 min), and the supernatant was transferred to a fresh low-protein-binding tube (Eppendorf). The beads were then washed with 2 × 250 μL HPLC-grade water, and the washes were combined with the original supernatant. The peptide solution was acidified with formic acid to a final concentration of 0.5%, and cleared from insoluble particulates by centrifugation for 10 min at 16,100 × *g* (4 °C), and the supernatant was transferred to clean tubes. Samples were dried in a SpeedVac concentrator and resuspended in 100 μL HPLC-grade water. Methionine oxidation was performed by the adding H_2_O_2_ to a final concentration of 0.5% and incubating for 30 minutes at 30 °C. Solid-phase extraction of peptides was performed using C18 reversed-phase sorbent pipette tips (Bond Elut OMIX 100 µL C18 tips, Agilent, Santa Clara, CA, USA) according to the manufacturer’s instructions. The tips were conditioned by aspirating the maximum pipette tip volume of water:acetonitrile (50:50, v/v), which was then discarded. After equilibration by washing 3 times with 0.1% TFA in water, 2 × 100 µL of the acidified sample were loaded and aspirated through the tip 10 times to maximize binding efficiency. The tip was washed 3 times with 0.1% TFA in water:acetonitrile (98:2, v/v), and bound peptides were eluted into LC-MS/MS using 0.1% TFA in water:acetonitrile (30:70, v/v). The samples were vacuum-dried in a SpeedVac concentrator and re-dissolved in 20 µL of 2 mM tris (2-carboxyethyl) phosphine in 2% acetonitrile.

### LC-MS/MS analysis

7.5 µL of each sample was injected for LC-MS/MS analysis on an Ultimate 3000 RSLC nano HPLC system (Thermo) in-line connected to a Q Exactive HF mass spectrometer (Thermo Fisher Scientific Inc.) equipped with a Nanospray Flex Ion source (Thermo), as previously described (74, 75). Specifically, trapping was performed at 10 μL/min for 4 min in solvent A on a 20 mm trapping column (made in-house, 100 μm internal diameter (I.D.), 5 μm beads, C18 Reprosil-HD, Dr. Maisch, Germany) and the sample was loaded on a 200 cm long micropillar array column (PharmaFluidics) with C18-end-capped functionality mounted in the Ultimate 3000’s column oven at 50 °C. For proper ionization, a fused silica PicoTip emitter (10 µm inner diameter; New Objective) was connected to the µPAC™ outlet union, and a grounded connection was provided to this union. Peptides were eluted by a non-linear increase from 1 to 55% MS solvent B (0.1% formic acid in water/ACN (2:8, v/v)) over 75 minutes, initially at a flow rate of 750 nL/min, then at 300 nL/min, followed by a 15-minute wash step reaching 99% MS solvent B and re-equilibration with MS solvent A (0.1% formic acid in water/ACN (2:8, v/v)). The mass spectrometer was operated in data-dependent, positive ionization mode, automatically switching between MS and MS/MS acquisition for the 16 most abundant ion peaks per MS spectrum. Full-scan MS spectra (*m*/*z* 375-1500) were acquired at a resolution of 60,000 in the Orbitrap analyser after accumulation to a target value (AGC target) of 3,000,000. The top 16 most intense ions above a threshold of 13,000 were isolated (1.5 Th window) for fragmentation using a normalised collision energy of 28%, after filling the trap to a target value of 100,000 for a maximum 80 ms. MS/MS spectra, with a fixed first mass of 145 *m*/*z*, were acquired at a resolution of 15,000 in the Orbitrap. The S-lens RF level was set to 50, and precursor ions with single, unassigned and >7 charge states were excluded from fragmentation. Raw data files were searched using MaxQuant version 1.6.10.43 (37). Spectra searched against the UniProt database restricted to *Solanum lycopersicum* (TaxID 4081, UP000004994, version 2022_01, containing 34,655 protein entries; https://www.uniprot.org/proteomes/UP000004994) to which the tagged bait sequences (i.e., TurboID tagged eGFP and Mj-MSP18_ΔSP_) were added. MaxQuant also automatically included potential contaminants from the *contaminants.fasta* file. Multiplicity was set to 1 (no labeling), and label-free quantitation (LFQ) was performed using MaxQuant’s default settings, without “match between runs”. Acetylation (protein N-term) was selected as a variable modification, and methionine oxidation (to methionine sulfoxide) was set as a fixed modification, since all methionines were uniformly oxidized using hydrogen peroxide.. The enzyme rule was set to trypsin/P, allowing a maximum of one missed cleavage. The main search peptide tolerance was 4.5 ppm, and the MS/MS ion trap match tolerance was 0.5 Da. The peptide-spectrum match (PSM) FDR was set at 1%, with a minimum Andromeda score of 40 for modified peptides. Protein-level FDR was also set at 1%, estimated by using reversed sequences. The maximum number of modifications per peptide was set to 5. For protein quantification (from the *proteinGroups.txt* file), only unique peptides were used, and all modifications were allowed. A multiscatter plot with Pearson correlation was generated in Perseus **(Figure S4)** for quality control across the biological replicates. Subsequent statistical analysis was performed using a two-sample *t*-test on imputed LFQ intensities to identify significantly enriched plant proteins in the effector samples compared to the GFP control (37, 38). Plotting and visualizations were done using Python 3.9 and Matplotlib.

### Phylogenetic analyses

The tomato BSK7 protein sequence was compared with 66 homologous proteins of the BSK family across *A. thaliana*, *N. benthamiana* and *S. lycopersicum* based on similarity to the conserved protein kinase domain. 113 Mj-MSP18 putative homologous proteins derived from *M. incognita, M. hapla*, *M. arenaria*, *M. enterolobii* and *M. graminicola* species were compared to generate their phylogenetic tree. Multiple sequence alignments were performed with Muscle (Edgar, 2004) as implemented in MEGA v11 (Edgar, 2004). Phylogenetic reconstructions were obtained by the NJ (neighbor-joining) method (Saitou & Nei, 1987) using the Jones-Taylor-Thornton (JTT) substitution model with Gamma-distributed rates (5 categories) among sites together with bootstrap analysis using 1000 replicates.

### Yeast two- or three-hybrid

Y2H and Y3H analysis was performed as described in (43, 76, 77), with the GAL4 system. Briefly, bait and prey open reading frames were fused to the GAL4-AD or GAL4-BD via cloning into pGADT7 or pGBKT7, respectively. The *S. cerevisiae* PJ69-4α and Y2HGold yeast strain was co-transformed with bait and prey using the polyethylene glycol (PEG)/lithium acetate method. Transformants were selected on Synthetic Defined (SD) media lacking leucine and tryptophan (Clontech). For Y2H three individual clones were grown ON in liquid cultures (lacking leucine and tryptophan) at 30 °C and 10x or 100x dilutions were spotted on the control media (SD-Leu-Trp) and selective media lacking Leu, Trp, and His (Clontech) with and without the addition of 3-amino-1H-1,2,4-triazole (Acros Organics (Thermo Fisher Scientific)) to test the strength of the interaction. For Y3H individual clones were grown ON in liquid cultures (lacking Leu, Trp and U) at 30°C. The individual colonies were spotted on the control media (SD-LWU) and selective media lacking Leu, Trp, U and His (Clontech) with and without the addition of 3-amino-1H-1,2,4-triazole (Acros Organics (Thermo Fisher Scientific)) to test the strength of the interaction. Additionally, the X-α-Gal (Clontech) assay was employed to quantitatively evaluate the intensity.

### Cell imaging and ratiometric BiFC

For microscopy, *A. tumefaciens* strain GV3101 transformed with the different constructs **(Supplemental Table 10)** was infiltrated into *N. benthamiana* leaves at OD600 final concentration of 0.3, then the tobacco plants were kept in daylight at room temperature (RT). In the subcellular localization assay, the leaf spots infiltrated 48 hours post-inoculation were imaged using a confocal laser-scanning microscope (Nikon Instruments Inc., Tokyo, Japan). eGFP was excited with a wavelength of 488 nm and emission was detected at 495–530 nm; eRFP was excited with a wavelength of 561 nm and emission was detected at 592–632 nm.

For rBiFC, *Sl-BSK7* and *Mj-MSP18_ΔSP_* were fused to the C-terminal fragment of mVenus (78) and the N-terminal fragment of mVenus, respectively, in the same 2 in 1 destination vector. Empty vector and the combination of non-interacting part of *Sl-BSK7(Sl-BSK7_Δ57-486_)* fused to *mVenusC* and *Mj-MSP18_ΔSP_* tagged with *mVenusN* in the same vector were both used as controls. Each construct was transformed into *A. tumefaciens* strain GV3101 for infiltration, and the mVenus signal was analyzed 72 h after infiltration into *N. benthamiana* leaves. mVenus was excited with a wavelength of 514 nm and emission was collected at 530–575 nm using confocal microscopy (Nikon Instruments Inc., Tokyo, Japan). Meanwhile, for the detection of RFP, the excitation and emission parameters were utilized as mentioned above. All the images were taken using the same setting, and the interacting pair was evaluated compared with the control based on the mVenus/RFP relative fluorescent intensity ratio. For statistical analysis, we considered not only the mean ratios obtained for each pair but also the impact of 7-9 biological cells. Each biological replicate included three technical replicates.

### Luminescence imaging

*Sl-BSK7* and *Sl-FlS2* were fused to the C-terminal and N-terminal fragments of firefly luciferase, respectively, in the binary vectors pCAMBIA1300::C-LUC and pCAMBIA1300::N-LUC (79). Split luciferase complementation assays of Sl-BSK7 and Sl-FlS2 were conducted by quantifying luminescence in five leaf discs sampled from agro-infiltrated regions of three plants, as previously described (27). The *Agrobacterium* carrying *Mj-MSP18_ΔSP_* or *eGFP* under control of the 35S promoter (vector pK7WG2, **Supplemental Table 10**) was grown for 2 days at 28 °C in LB medium with 25 μg/mL rifampicin, 25 μg/mL gentamicin, or 100 μg/mL spectinomycin and subsequently co-infiltrated with the *Agrobacterium* transformed with the obtained binary vectors into *N. benthamiana* leaves. Besides, empty binary vectors and the co-expression of Sl-BSK7-cLUC and Sl-FLS2-nLUC serve as additional controls. To assess luciferase activity in presence of MSP18_ΔSP_ and GFP *in planta*, tobacco leaves were treated with a 1 mM D-Luciferin solution (Duchefa; dissolved in 0.01% [v/v] Tween80, 0.1% [v/v] DMSO) 48 hours post agro-infiltration. The samples were then kept in total darkness for 5 minutes before imaging. The NightShade LB 985 (Berthold) system was used to capture the luminescence signal. The signal emitted by the sample was finally accumulated in the Charge-coupled device camera built-in the Nightshade system for 5 min, in total darkness. The images were acquired using a 1024 × 1024 pixels resolution (2 × 2 binning) with the IndiGO™ software. For analysis purpose, the images were first exported from the IndiGO™ software using the movie composer tool and choosing minimum and maximum threshold values by default, then the resulting files were imported in ImageJ then converted to 8-bit images for quantifying the signal intensity. The results were analyzed using four independent biological and three technical replicates.

### ROS production assay in *N. benthamiana* leaves

The *Agrobacterium* carrying *Mj-MSP18_ΔSP_* under control of the 35S promoter (vector pK7WG2, **Supplemental Table 10**) was grown for 2 days at 28 °C in LB medium with 25 μg/mL rifampicin, 25 μg/mL gentamicin, or 100 μg/mL spectinomycin. A plasmid expressing eGFP was used as a negative control; the aphid (*Myzus persicae*) effector Mp10 was used as a positive control (80). Flg22 (cat. no. RP19986, GenScript) was used as an inducer of ROS production. The protocol was performed as described before (81). Briefly, before infiltration, the overnight *Agrobacterium* cultures were washed and diluted in infiltration buffer (10 mM MgCl_2_,10 mM MES and 0.2 mM acetosyringone), to reach OD600 of 0.3 and incubated for three hours in the dark at room temperature. The next day, *N. benthamiana* 5–6 weeks old leaves were infiltrated with the *Agrobacterium* using a needleless syringe. The appropriate *Agrobacterium* was infiltrated at different spots on the same leaf. For each assay, one leaf from eight plants was used with two replicates per spot (each treatment contained eight biological and three technical replicates). Approximately 30 h after infiltration, 16 mm^2^ leaf discs were collected from the infiltrated areas with a cork borer, transferred to 96-well plates, and floated overnight on 200 μL filter-sterilized ultrapure water for recovery. After 48 h, the water was removed and replaced by a mixture of 100 nm flg22 (QRLSSGLRINSAKDDAAGLAIS (82)), 0.5 mM luminol probe 8-amino-5-chloro-7-phenyl-pyrido[3,4-d] pyridazine-1,4(2H,3H) dione (L-012) (Wako Chemicals, Richmond, Virginia, USA) and 20 μg/mL horseradish peroxidase (Sigma, Saint Louis, Missouri, USA). ROS production was measured by a luminol-based assay (83) over 60 min with integration at 750 ms.

### Cell death assay

*A. tumefaciens* strain GV3101 was used for relevant constructs (see **Supplemental Table 10**); plasmid pK7WG2-GFP was used as a negative control. Agrobacteria carrying a plasmid for expression of *Mj-MSP18_ΔSP_* or *INF1* (84) were grown for 2 days in 10 mL of LB medium with the appropriate antibiotics. Before infiltration, the cells were pelleted, washed, resuspended in infiltration buffer (10 mM MgCl_2_,10 mM MES and 0.2 mM acetosyringone), and incubated for at least 3 h at RT. Depending on the combination of constructs, the final concentration in the mixtures was adjusted to an OD600 of 0.5. The mixtures were spot-infiltrated into *N. benthamiana* leaves of 5 to 6-week-old plants. The negative controls were infiltrated on the same leaf as the tested effector. For each plant, two leaves were infiltrated, and 20 plants were used per assay. The response was recorded two days post infiltration. HR on the spot was noted as being suppressed when less than 50% of that spot showed cell death following (85). Fisher’s exact test was used to analyse the results statistically.

### Callose deposition assay

Callose deposits were observed according to the protocol with small modifications (86, 87). Briefly, *Mj-MSP18_ΔSP_* or *eGFP* (**Supplemental Table 10** for the vectors used) were infiltrated into *N. benthamiana* leaves using Agrobacterium GV3101 of an OD600 of 0.5. Accordingly, 48 h post infiltration the leaves were treated with 500 nM flg22 for 24 h. After that, they were fixed and destained in absolute ethyl alcohol: acetic acid, 3:1 v/v for 24 h followed by 1 h 70% ethanol, 1 h 50% ethanol 1 h 10% NaOH at 37 °C. Transparent leaf segments were washed three times in 150 mM K_2_HPO_4_ (pH 9.6) and stained with 0.02 % aniline blue (cat.no: B8563-500ML; Sigma) in 150 mM K_2_HPO_4_ (pH 9.6). The pictures were taken using a wide field epifluorescent microscope (Nikon Instruments Inc., Tokyo, Japan). Images were acquired with DAPI filter (excitation at 387/11 nm and emission 447/60 nm). Callose deposits were analyzed in fields of 8.4 mm^2^ using ImageJ (88) software and normalized to 1 mm^2^. Three technical and six biological repeats were used.

### Plant preparation and infection assays

*Meloidogyne javanica* eggs were harvested from infected tomato roots, sterilized with NaOCl, and hatched in a sodium azide solution followed by antibiotic treatment for surface sterilization. The hatching and sterilization process was adapted from previous methodologies (89, 90). Ten weeks post-inoculation, the roots were rinsed under running water to remove sand particles and then incubated in 0.05% (v/v) NaOCl with continuous shaking for 3 min, followed by sieving (Hussey & Barker, 1973). The eggs were collected using a 25 µm sieve. Subsequently, the eggs were incubated in 0.02% sodium azide (NaN_3_) for 20 min while shaking. Afterward, the eggs were thoroughly rinsed with tap water to remove traces of NaN_3_ and incubated in a 25 µm sieve in a solution of 1.5 mg/mL gentamycin and 0.05 mg/mL nystatin in the dark at room temperature. After four days, the hatched second-stage juveniles (J2s) were collected, purified by separation in a 70% sucrose column, and then surface sterilized with 0.16 mM HgCl_2_, 0.49 mM NaN_3_, and 0.002% (v/v) Triton-X for 20 min. The J2s were rinsed three times with sterile tap water and placed in a 0.07% (w/v) Gelrite solution (Duchefa Biochemie, Haarlem, the Netherlands). *A. thaliana* seeds wild-type Col-0 and *fls2* mutants, *bsk7, bsk8* single mutants and *bsk7,8* double mutants and *S. lycopersicum* seeds *Sl-bsk7 (fir1)* and *Sl-fls2* mutants, along with their respective wild-type plants, Hawaii 7981 and Rio Grande prf3 obtained as previously described (25, 27) were sterilized, germinated, and grown under controlled conditions before inoculation with *M. javanica* juveniles. Gall and egg mass counts were obtained from three independent experiments (n ≥ 29 per genotype), with data normalization to account for batch effects as described previously (91). Statistical analyses were performed using Wilcoxon rank’s test to assess differences in susceptibility among genotypes, utilizing Rstudio version 4.0.2×64 (R Core Team 2023) for data management and visualization.

### RNA-seq sample preparation

Seeds of tomato (*S. lycopersicum* cv Moneymaker) were surface sterilized in 70% (v/v) ethanol for 10 min and then rinsed three times 5 min with sterile water. The seeds were germinated on MS tissue culture medium containing 4.3 g/l MS medium (Duchefa, catalog no. M0221.0050), 0.5 g/L MES, 30 g/L sucrose, pH 5.8, and 10 g/L agar (Difco, catalog No. 214530) in dark. Seeds were germinated at 24 °C in a growth chamber (16-h-light/8-h-dark photoperiod) for 10–14 days until cotyledons were fully expanded and the true leaves just emerged. *R. rhizogenes* strain ATCC 15834 transformation was performed as described previously by (92) with some minor modifications. More specifically, competent rhizogenic Agrobacterium cells were transformed by electroporation (93) with the desired binary vector, plated on yeast extract beef (YEB) medium plates with the appropriate antibiotics (100 mg/L spectinomycin), and incubated for 3 to 4 days at 28 °C. A transformed Agrobacterium culture was inoculated from fresh plates into YEB liquid medium with the appropriate antibiotics added and grown overnight at 28 °C with shaking at 200 rpm. The cut cotyledons were soaked in liquid bacterial suspension at an OD at 600 nm of 0.2–0.4 in MS liquid medium for 20 min and transferred onto MS agar plates without antibiotics for 3 days. After that, the cotyledons were transferred to MS agar plates with 200 mg/L cefotaxime (Duchefa, catalog no. c0111.0025) and 50 mg/L kanamycin and returned to 24 °C. The screening for expression of the eGFP marker of antibiotic-resistant roots was monitored by using a wide-field epifluorescent microscope (Nikon Instruments Inc., Tokyo, Japan). Four independent roots showing expression of the marker were subcloned for each construct. After three rounds of cultivation, root cultures were maintained and grown in antibiotic-free MS medium supplemented with 3% sucrose (w/v) at 24 °C. Two weeks after growing in liquid medium, effector expression was induced with 100 µM β-estradiol (cat. no: E8875-1G; Sigma) for 24 h. After that samples were grounded and from one part RNA was extracted, while from the second half proteins were extracted and analyzed by western blot. For mRNA sequencing, RNA was extracted from ±100 mg ground tissue using the RNeasy Plant MiniKit (Qiagen). Four biological replicates, each independently transformed, were used per treatment. RNA quality control was performed. RNA 6000 nanochip (Agilent technologies) libraries were prepared using QuantSeq 3’ mRNA library prep FWD kit (Lexogen) according to the manufacturer’s instructions. Library-quality was verified using a High sensitivity DNA chip (Agilent technologies). Library quantification was assessed using qPCR. According to the Illumina protocol, equimolar pooling of libraries based on qPCR was done, and libraries were sequenced on an Illumina NextSeq 500 platform, SR76, with high output. The sequencing depth was 8M.

### RNA-seq data analysis

Data analyses were carried out using the Galaxy Australia platform (https://usegalaxy.org.au/) (94). The quality was assessed using FastQC (95) and MultiQC (96), and reads were trimmed with Trimmomatic (97). Performing the initial ILLUMINACLIP step (3,30,10,8) and two additional Trimmomatic steps: SLIDINGWINDOW (3,10) and MINLEN, reads < 20 bp were removed. Trimmed reads were mapped to the tomato reference genome (ITAG 3.0, (98)) using HISAT2 (99). Reads were counted using StringTie (Pertea et al., 2015). Differentially expressed genes were identified using DeSeq2 (100). Genes were considered differentially expressed if P adjusted < 0.05 and log2FC < 0 (downregulated) or log2FC > 0 (upregulated). The data were prepared using custom scripts https://github.com/borisstojilkovic/postdeseq2-annotate-and-split-deseq2-to-Goseq-files. GoSeq (101) and g: Profiler (102) were used for gene ontology analysis and pathway analysis (https://github.com/borisstojilkovic/g-Profiler-Batch-Enrichment-GO-terms-Python). Genome and annotation files were assessed from EnsemblPlants (https://plants.ensembl.org/index.html) (103). Genes belonging to each GO category were extracted using a custom Python script available at https://github.com/borisstojilkovic/gene-per-go-extractor. Plotting and visualizations were done using Python 3.9 and Matplotlib.

### qRT-PCR for confirmation of RNA-seq results

RNA was extracted using the RNeasy Plant Mini Kit (Qiagen), and cDNA was synthesized using Maxima First Strand cDNA Synthesis Kit for qRT-PCR (Thermo Fisher Scientific). Primers were designed using QuantPrime (104). qRT-PCR wasperformed with four biological replicates. The qPCR conditions consisted of initial denaturation at 95 °C for 10 min followed by 50 cycles of 95 °C for 25 s, 58 °C for 25 s, and 72 °C for 20 s (all primers were designed using QuantPrime (104) and are listed in the **Supplemental Table 11**). Expression data were normalized using three reference genes (*ARD, UBE2* and *DEAD40*) (105) and analyzed using REST2009 (106).

### cDNA library preparation

The cDNA library was already available from Alain Goossens lab. To generate this library, MoneyMaker plants were grown hydroponically for 4 weeks. Next, the plants were treated with either 500 µM salicylic acid, 50 µM jasmonate and OD0.1 Agrobacterium K599. Hormone-treated roots were harvested on different timepoints 2hrs, 1 day and in case of SA also 2 days post-treatment. Agro-treated roots were harvested 2 days, 6 days and 8 days post-treatment. RNA was extracted using the RNA Maxiprep Kit (Qiagen) following the manufacturer’s instructions. Further mRNA was isolated using the Poly(A)Purist™ MAG Kit (ThermoFisher) and subsequently the cDNA library was constructed using the CloneMiner™ II cDNA Library Construction Kit (ThermoFisher). The number of primary clones contained 1.39 x 108 cfu with an average insert size of 1.3 kb.

### Yeast cDNA library screen

Y2H-screen was performed as described previously (43). Briefly, bait proteins were first transformed into the *S. cerevisiae* PJ69-4A yeast strain. Competent bait yeast strains were super-transformed using 50 mg of the cDNA library made as described above and plated on SD-LW to determine the transformation efficiency and on SD-LWH+3-AT for identifying protein interactions. A transformation efficiency of at least 1 × 10^6^ clones was achieved.

### Motif analysis for MYB/MYC2 TFs binding site in promoter regions

First, promoter regions were extracted using extract_promoters_script.py, and then extracted sequences were used to perform motif search using motif_check_script.py. The scripts are using Python 3.9 and libraries (SeqIO from BioPython, Pandas, SciPy for statistics and Matplotlib for visualization). The first script (extract_promoters_script.py) extracts upstream and /or downstream region of the gene based on annotation (BED) and genome (FASTA) files giving as output a new fasta file with extracted promoters for a given set of genes. The second script uses given motifs to search in the extracted promoters file. The script also extracts genes containing the motif and calculates percentage of the genes containing the binding site and compares to all extracted promoters and calculates the p-value using Fisher’s exact test. All the code and detailed explanations are available on GitHub (https://github.com/borisstojilkovic/Extract-promoters-and-motif-scan).

### Extraction and quantification of plant hormones in tomato hairy roots

Tomato hairy root samples (approximately 10 mg fresh weight) were extracted in ice-cold 60% acetonitrile for 12 h at 4 °C and 25 pmol of deuterium-labelled internal standards of BRs was added to each sample (OlChemIm Ltd., Olomouc, Czech Republic). Then the samples were centrifuged (36 670 *g*, 15 min, 4 °C) and supernatants were passed through 50 mg Discovery™ DPA-6S cartridges (Supelco®, Bellefonte, PA, USA). After evaporation to dryness, samples were reconstructed in 40 µL of methanol and analyzed by liquid chromatography with tandem mass spectrometry (UHPLC-MS/MS) using an ACQUITY UPLC® I-Class System (Waters, Milford, MA, USA) combined with a triple quadrupole mass spectrometer Xevo™ TQ-S MS (Waters MS Technologies, Manchester, UK). The UHPLC-MS/MS analysis is described in (107) and (108). All data were processed using MassLynx software (ver. 4.1, Waters) and the analyses were performed in 4 repetitions. For determination of other plant hormones including jasmonic acid (JA), (-)-jasmonyl-L-isoleucine (JA-Ile), abscisic acid (ABA), and salicylic acid (SA), the hairy root samples (approximately 10 mg FW) were extracted in cold 1 mL of 1 mol/L formic acid in 10% aqueous methanol with the addition of internal standards labeled with stable isotopes (^2^H_6_)JA, (^2^H_2_)(-)-JA-Ile, (^2^H_6_)ABA and (^2^H_4_)SA according to (109) and the whole extracts were purified on reverse phase Oasis® HLB solid phase extraction columns (1cc/30 mg, Waters). The analysis was performed with 4 repetitions on an Agilent 6490 Triple Quadrupole LC/MS system coupled to a 1290 Infinity LC system (Agilent Technologies, Santa Clara, CA, USA) as described (109).

## Funding

This work was supported by the Special Research Fund of Ghent University (projects BOF18/GOA/013 and BOF23/CDV/032), and by the Research Foundation – Flanders (FWO-Vlaanderen, project number G045921N) awarded to P.V.D., the Czech Science Foundation (grant No. 22-17435S) and Chinese Scholarship Council doctoral fellowship awarded to Y.C.

## Acknowledgments

We extend our gratitude to Maria Aparicio Chacon and Sofie Goormachtig (Goormachtig Lab, VIB Center for Plant Systems Biology, Ghent, Belgium) for providing plasmids essential for localization studies and for their invaluable discussions. We also thank Lore Gryffroy for helping with the TurboID experiments. Guy Sobol for providing Arabidopsis and tomato mutants and sharing preliminary data.

## Competing interests

The authors declare no competing interests.

## Data and Software Availability

The mass spectrometry proteomics data generated in this study have been deposited to the ProteomeXchange Consortium via the PRIDE (110) partner repository with the dataset identifier PXD067607. During peer review, the dataset can be accessed using the reviewer login credentials:

Username

Password: SZXIzVJRERAv

The mRNA-sequencing data have been deposited in the NCBI Sequence Read Archive (SRA) under BioProject ID PRJNA1302155. All other data supporting the findings of this study are available within the manuscript and supporting information.

## Notes

### Competing Interest Statement

The authors have declared no competing interest.

