## Supplementary material for "The root-knot nematode effector Mj-MSP18: a Swiss army knife for reprogramming plant immunity": Figure S

### Supplemental figures and tables

#### Supplemental figures

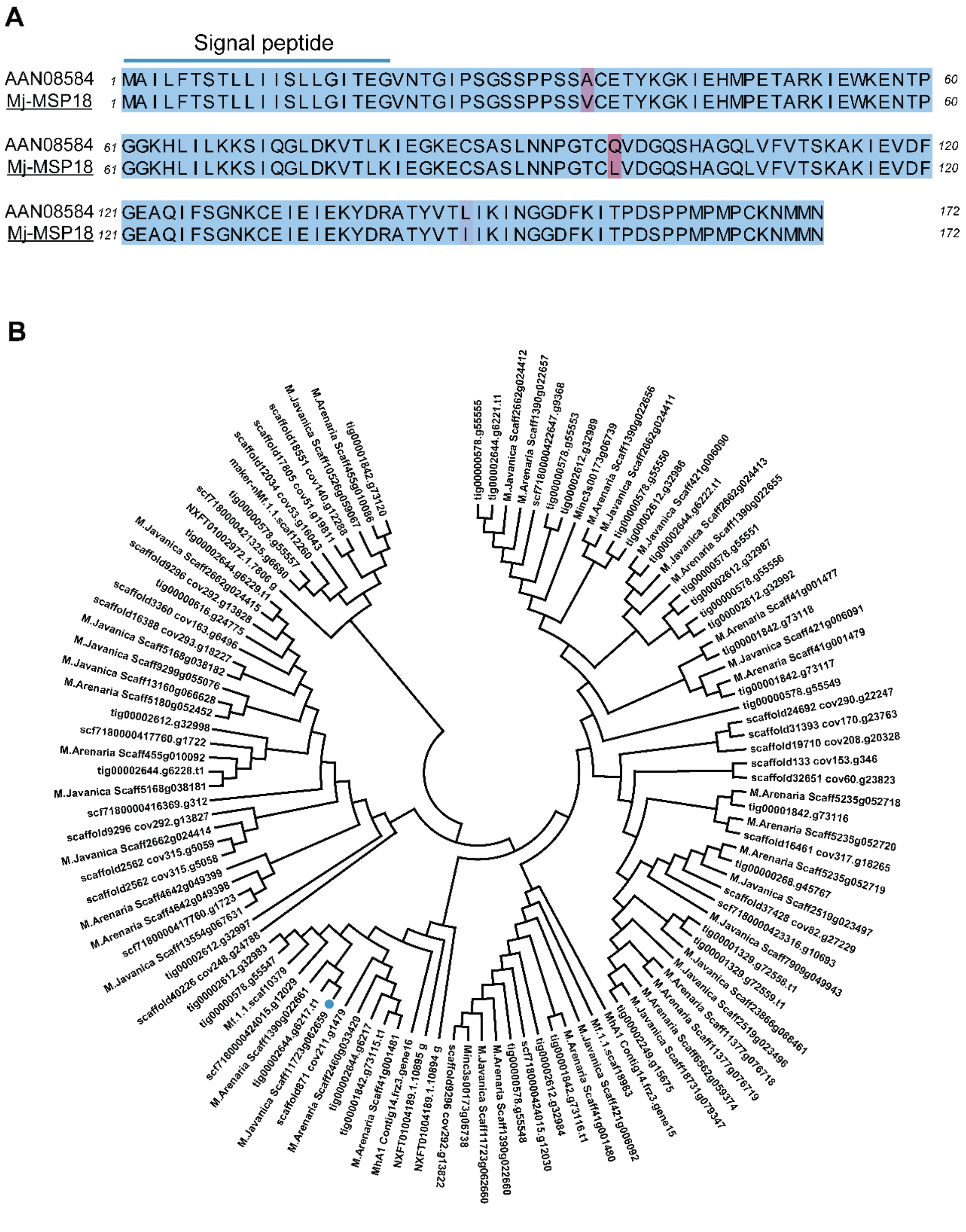


Fig. S1

**Figure S1. Homology analysis of Mj-MSP18**

(A) Alignment of Mj-MSP18 and its closest homolog from *M. incognita*. Alignment was performed with Clustal Omega and visualized via Jalview. AAN08584 is NCBI accession number of *M. incognita* putative oesophagal gland cell secretory protein 18. Blue color indicates identical amino acids between the two protein sequences while red color shows distinct physicochemical properties of amino acids at specific positions. The darker the red, the greater the disparity in the chemical properties.

(B) Phylogenetic tree of Mj-MSP18 homologous proteins across various *Meloidogyne* species. Sequences are derived from the WormBase ParaSite and INRAE database and encompassing *M. incognita, M. hapla*, *M. arenaria*, *M. enterolobii* and *M. graminicola* species. The classification was executed using Clustal Omega and visualized with MEGA_11. The blue dot indicates Mj-MSP18.

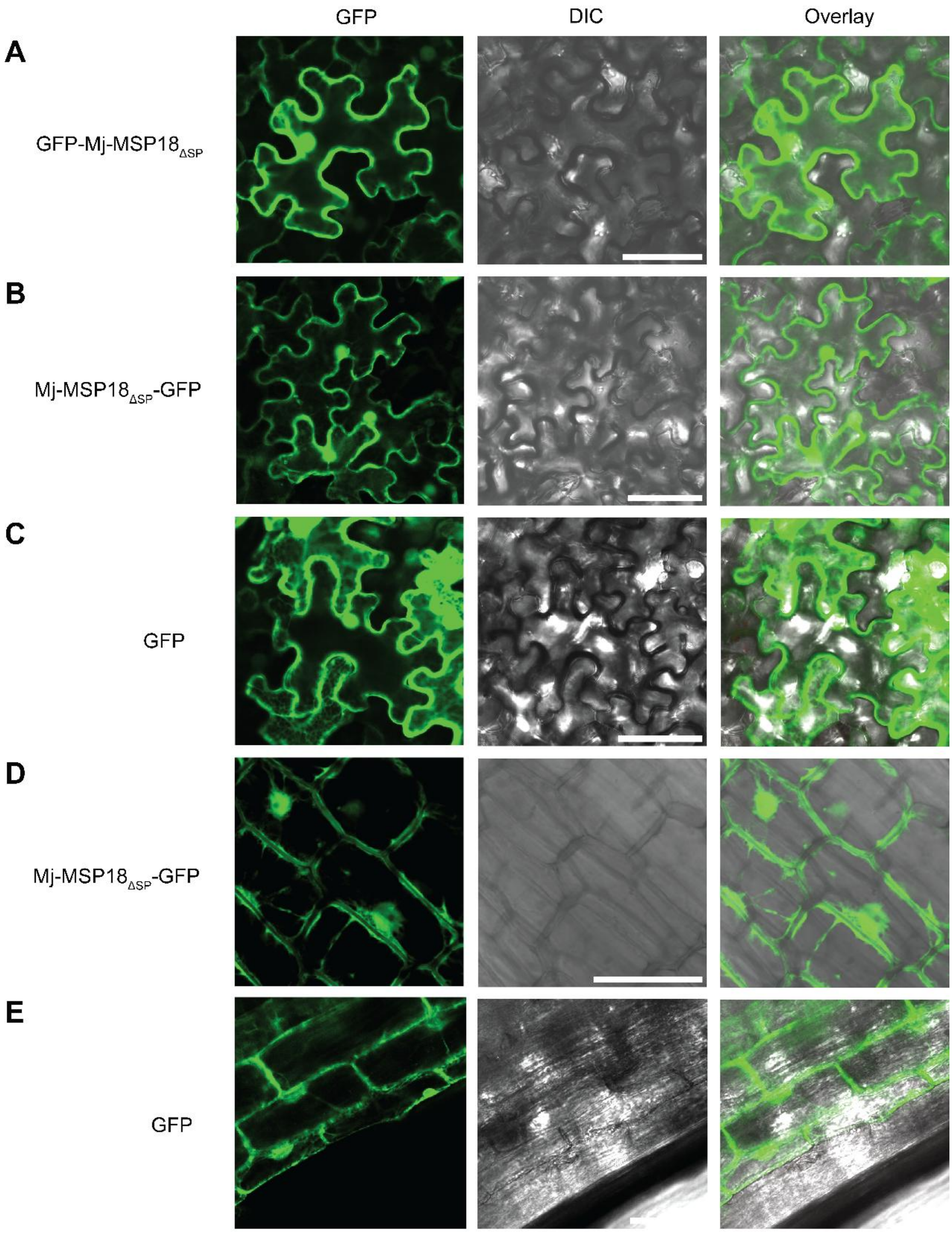
GFP
DIC
Overlay
A
GFP-Mj-MSP18ΔSP
B
Mj-MSP18ΔSP-GFP
C
GFP
D
Mj-MSP18ΔSP-GFP
E
GFP

Fig. S2

**Figure S2. Subcellular localization of Mj-MSP18$_{\Delta SP}$ and GFP in *N. benthamiana* and tomato hairy roots**

*GFP* was fused to Mj-*MSP18$_{\Delta SP}$* at the N-terminus (A) and C-terminus (B). These fusion constructs and individual *GFP* (C), were transiently expressed under the control of the constitutive CaMV35S promoter in *N. benthamiana* leaves. The *GFP* fluorescent reporter gene C-terminally fused to the *Mj-MSP18$_{\Delta SP}$* was also transformed in tomato hairy roots under the control of the constitutive CaMV35S promoter (D) while individual *GFP* (E) was expressed under control of prolD (pKCTAP). Observation was done by fluorescence microscopy (Nikon Instruments Inc., Tokyo, Japan). Scale bars = 50 µm.

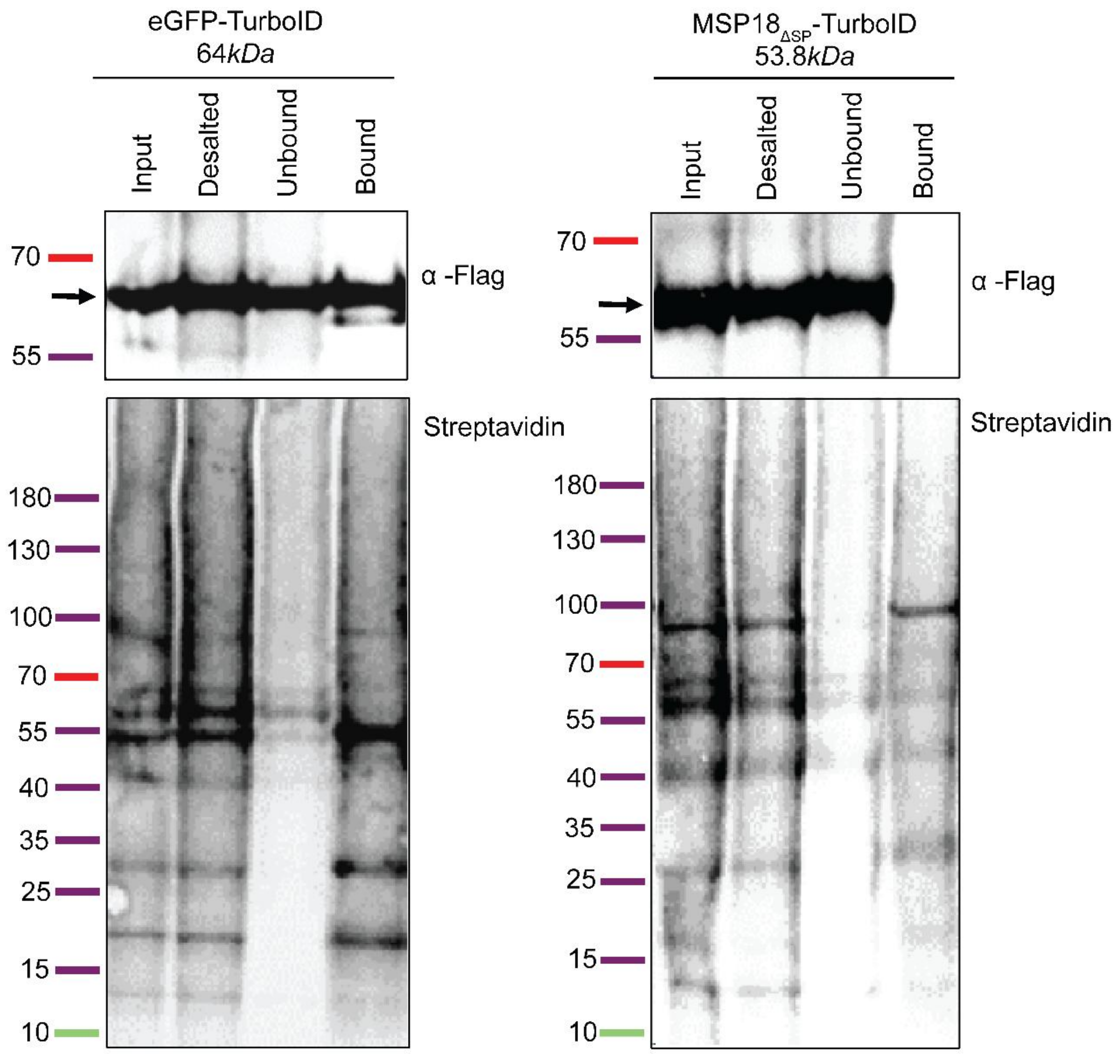
eGFP-TurboID
64kDa
Input
Desalted
Unbound
Bound
70
55
α -Flag
Streptavidin
180
130
100
70
55
40
35
25
15
10
MSP18ΔSP-TurboID
53.8kDa
Input
Desalted
Unbound
Bound
70
55
α -Flag
Streptavidin
180
130
100
70
55
40
35
25
15
10

Fig. S3

**Figure S3. Intactness and biotinylation of the TurboID-fusion proteins in tomato hairy roots.**

XVE::eGFP-TurboID and XVE::MSP18$_{\Delta SP}$-TurboID constructs were used for rhizogenic *Agrobacterium*-mediated transformation of tomato hairy roots. The XVE system is based on a chimeric transcription factor that combines several important functional domains, specifically: "X" refers to the LexA DNA-binding domain, "V" stands for the VP16 transactivating domain, and "E" represents the estrogen receptor (ER) regulatory region. Transformed roots were treated with 100 μM β-estradiol for 22-hour induction, followed by a 2-hour supplementation with 50 μM biotin. Prior to subjecting protein extracts from hairy roots to mass spectrometry, the quality of the TurboID-MS sample preparation was assessed by immunoblot of the input, input desalted, unbound and bound fractions from the eGFP- and Mj-MSP18$_{\Delta SP}$-TurboID-fusion proteins. The anti-Flag antibody was utilized to detect the translational fusion of eGFP- and Mj-MSP18$_{\Delta SP}$-TurboID-Flag (top panels) and a streptavidin/Alexa Fluor™ 680 conjugate was employed for visualizing their biotinylation patterns (bottom panels). Arrows indicate the size of fusion proteins eGFP (top left) and MSP18$_{\Delta SP}$ (top right). The results shown are representative of the three biological TurboID replicate hairy root samples.

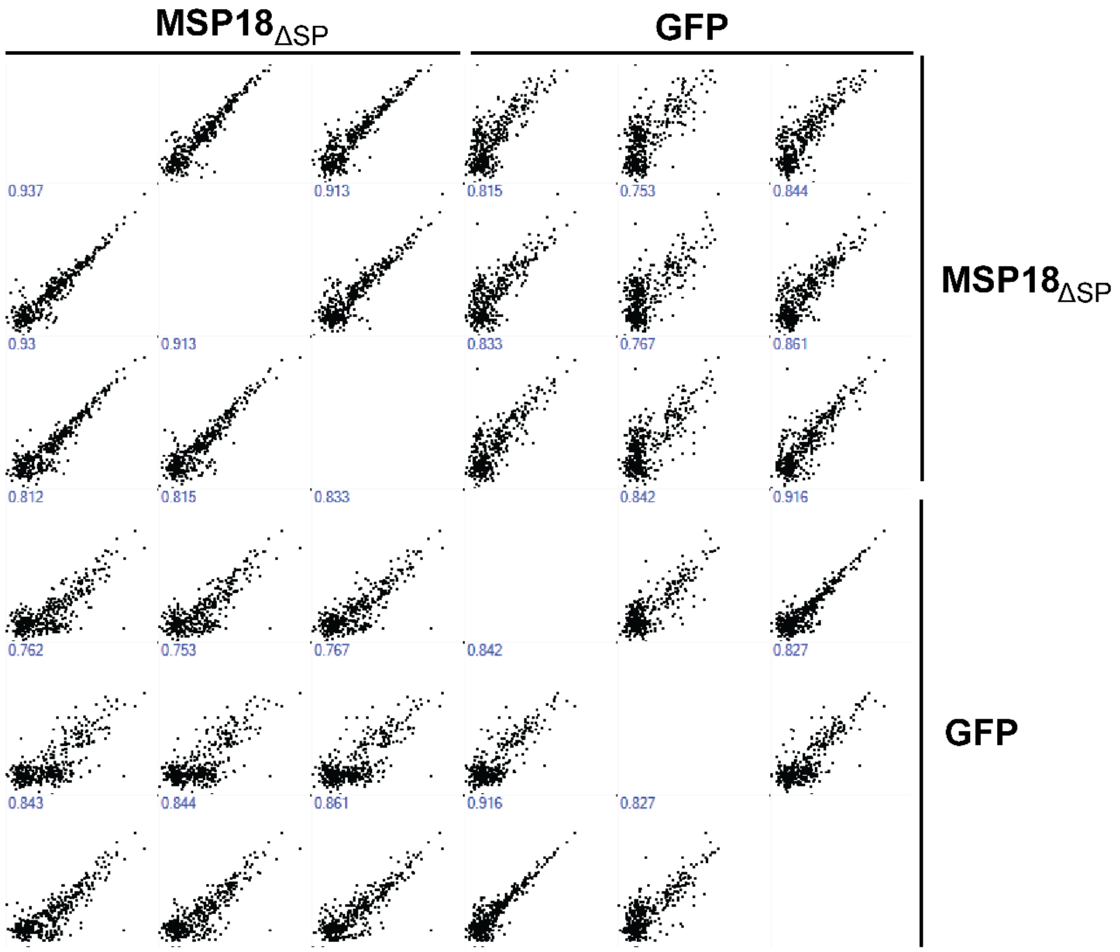

MSP18ΔSP
GFP
0.937
0.913
0.815
0.753
0.844
0.93
0.913
0.833
0.767
0.861
0.812
0.815
0.833
0.842
0.916
0.762
0.753
0.767
0.842
0.827
0.843
0.844
0.861
0.916
0.827
MSP18ΔSP
GFP


Fig. S4

**Figure S4. Multiscatter plots of the Mj-MSP18$_{\Delta SP}$ and GFP TurboID screens.** Multiscatter plots with Pearson correlations show the quality and reproducibility between the different samples and biological replicates in the Mj-MSP18ΔSP and GFP TurboID screens.

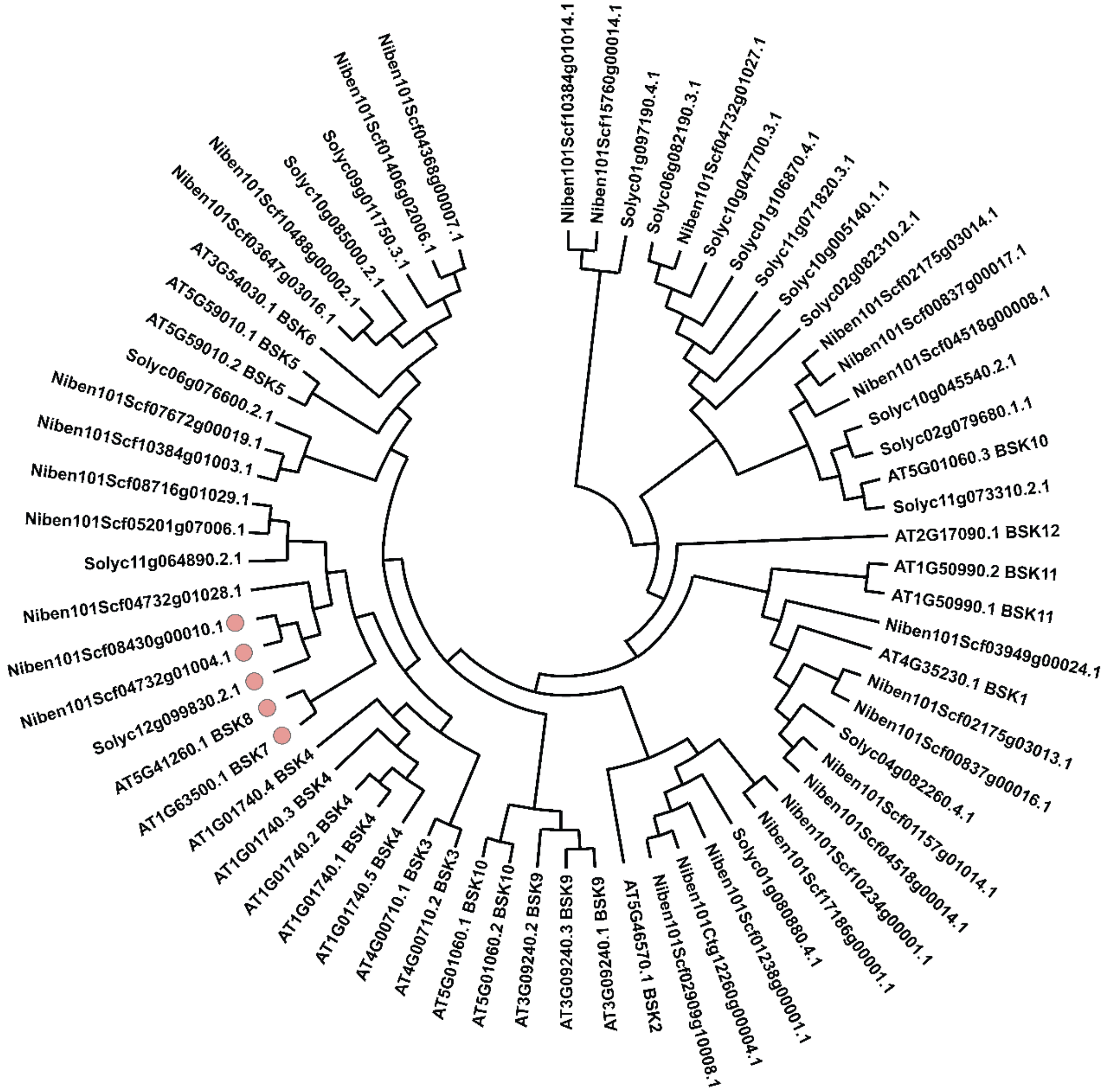
Niben101Scf10384g01014.1
Niben101Scf15760g00014.1
Solyc01g097190.4.1
Solyc06g082190.3.1
Niben101Scf04732g01027.1
Solyc10g047700.3.1
Solyc01g106870.4.1
Solyc11g071820.3.1
Solyc10g005140.1.1
Solyc02g082310.2.1
Niben101Scf02175g03014.1
Niben101Scf00837g00017.1
Niben101Scf04518g00008.1
Solyc10g045540.2.1
Solyc02g079680.1.1
AT5G01060.3 BSK10
Solyc11g073310.2.1
AT2G17090.1 BSK12
AT1G50990.2 BSK11
AT1G50990.1 BSK11
Niben101Scf03949g00024.1
AT4G35230.1 BSK1
Niben101Scf02175g03013.1
Niben101Scf00837g00016.1
Solyc04g082260.4.1
Niben101Scf01157g01014.1
Niben101Scf04518g00014.1
Niben101Scf10234g00001.1
Niben101Scf17186g00001.1
Solyc01g080880.4.1
Niben101Scf01238g00001.1
Niben101Ctg12260g00004.1
Niben101Scf02909g10008.1
AT5G46570.1 BSK2
AT3G09240.1 BSK9
AT3G09240.3 BSK9
AT3G09240.2 BSK9
AT5G01060.2 BSK10
AT5G01060.1 BSK10
AT4G00710.2 BSK3
AT4G00710.1 BSK3
AT1G01740.5 BSK4
AT1G01740.1 BSK4
AT1G01740.2 BSK4
AT1G01740.3 BSK4
AT1G01740.4 BSK4
AT1G63500.1 BSK7
AT5G41260.1 BSK8
Solyc12g099830.2.1
Niben101Scf04732g01004.1
Niben101Scf08430g00010.1
Niben101Scf04732g01028.1
Solyc11g064890.2.1
Niben101Scf05201g07006.1
Niben101Scf08716g01029.1
Niben101Scf10384g01003.1
Niben101Scf07672g00019.1
Solyc06g076600.2.1
AT5G59010.2 BSK5
AT5G59010.1 BSK5
AT3G54030.1 BSK6
Niben101Scf03647g03016.1
Niben101Scf10488g00002.1
Solyc10g085000.2.1
Solyc09g011750.3.1
Niben101Scf01406g02006.1
Niben101Scf04368g00007.1

Fig. S5

**Figure S5. Protein phylogenetic tree of BSK family.**

Genes of BSK across *A. thaliana*, *N. benthamiana* and *S. lycopersicum* are clustered based on similarity to the conserved protein kinase domain with Clustal Omega. Pink dots represent Sl-BSK7, two homologous proteins to Sl-BSK7 from *N. benthamiana* and another two BSK7 homologous proteins in *A. thaliana*.

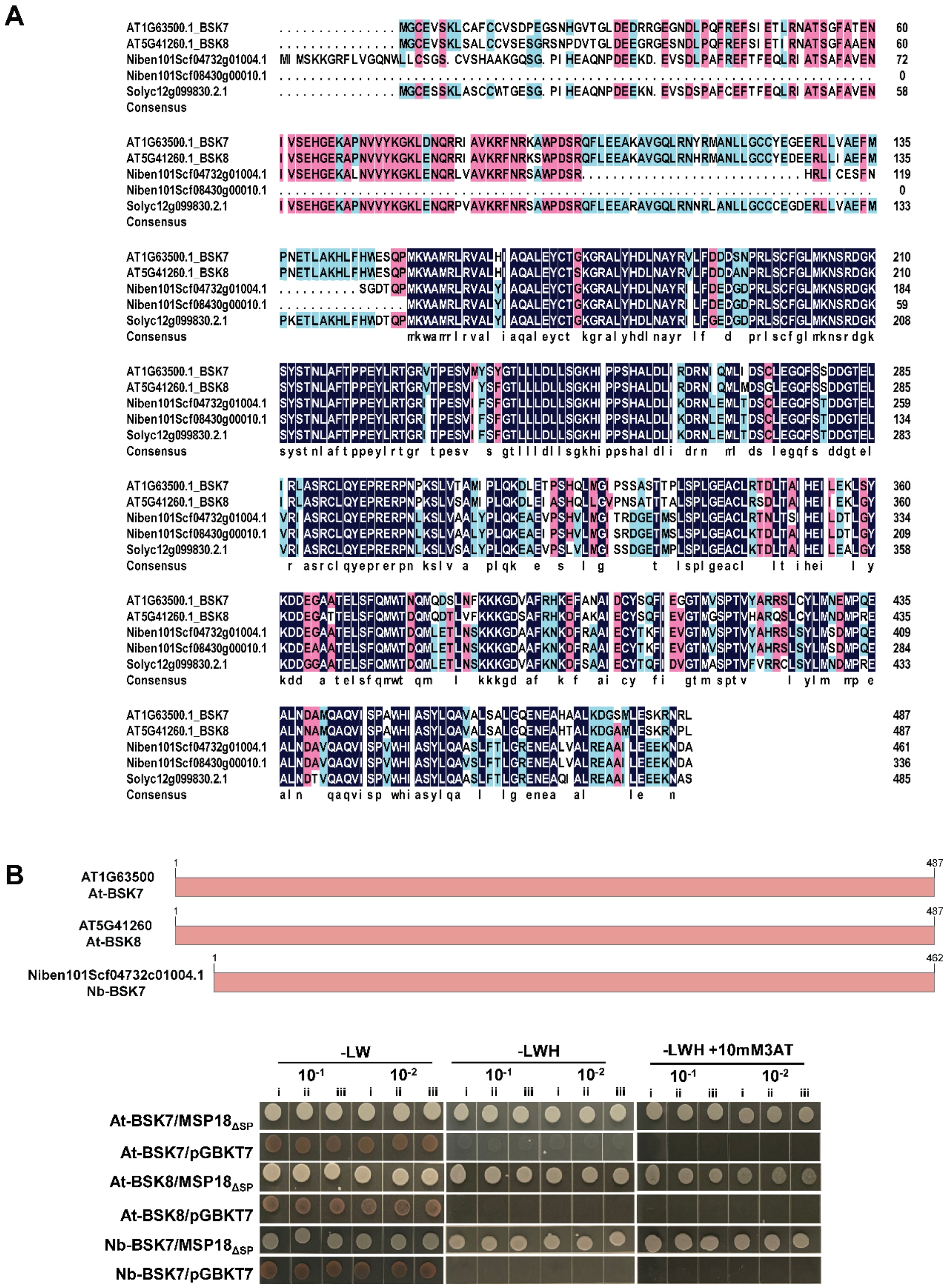
A
AT1G63500.1_BSK7
AT5G41260.1_BSK8
Niben101Scf04732g01004.1
Niben101Scf08430g00010.1
Solyc12g099830.2.1
Consensus
B
AT1G63500
At-BSK7
AT5G41260
At-BSK8
Niben101Scf04732c01004.1
Nb-BSK7
1
487
462
-LW
-LWH
-LWH +10mM3AT
$10^{-1}$
$10^{-2}$
i ii iii
At-BSK7/MSP18$_{\Delta SP}$
At-BSK7/pGBKT7
At-BSK8/MSP18$_{\Delta SP}$
At-BSK8/pGBKT7
Nb-BSK7/MSP18$_{\Delta SP}$
Nb-BSK7/pGBKT7

Fig. S6

**Figure S6. BSK7 homologous protein alignment and interactions between Mj-MSP18$_{\Delta SP}$ and BSK7 homologs of different plant species.**

(A) Alignment of the five closest BSK7 homologous proteins from *A. thaliana*, *N. benthamiana* and *S. lycopersicum* protein database in DNAMAN. The degree of similarity of the protein sequences is shown in distinct colors (50–74% cyan; 75–99%, cherry red; deep blue, 100%).

(B) Investigating the interactions between Mj-MSP18$_{\Delta SP}$ and three Sl-BSK7 homologous proteins of comparable length derived from *A. thaliana* and *N. benthamiana* by yeast two-hybrid assay. Schematics of the tested *A. thaliana* and *N. benthamiana* BSK7 protein are represented in the upper panel. Yeast transformants expressing Mj-MSP18$_{\Delta SP}$ were evaluated for their growth on SD-LW or SD-LWH with or without additional 3AT (3-amino-1,2,4-triazole), in combination with At-BSK7 (AT1G63500), At-BSK8 (AT5G41260), or Nb-BSK7 (Niben101Scf04732c01004). For the autoactivation assay, the same yeast strain was transformed with constructs encoding the At-BSK7, At-BSK8 or Nb-BSK7 proteins individually, and a complementary empty vector on the same medium. The tested truncated protein versions are mentioned on the left. Dilutions are indicated on top; i, ii and iii indicate different biological replicates.

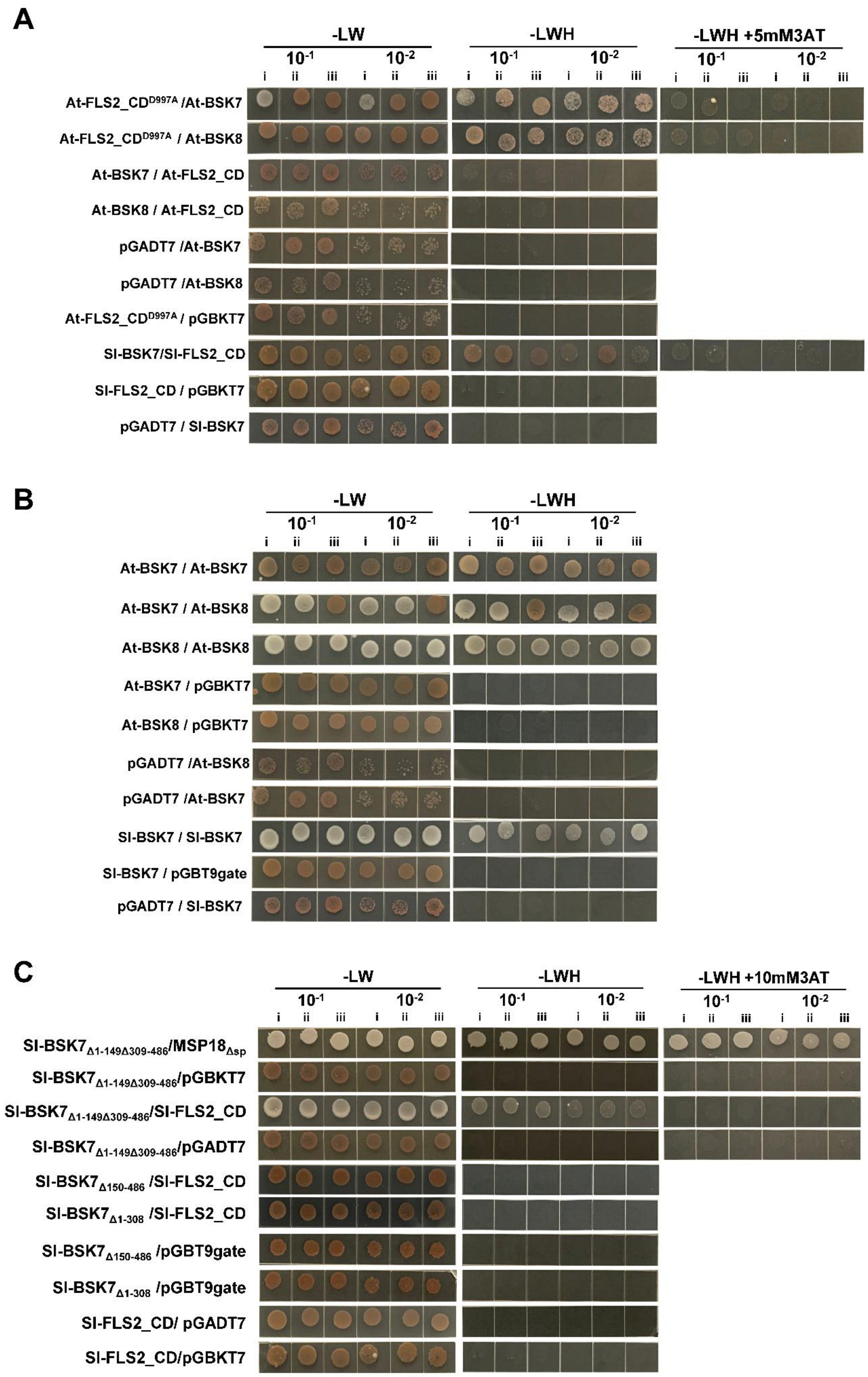
A
-LW
-LWH
-LWH +5mM3AT
$10^{-1}$
$10^{-2}$
i ii iii i ii iii
At-FLS2_CD$^{D997A}$ /At-BSK7
At-FLS2_CD$^{D997A}$ / At-BSK8
At-BSK7 / At-FLS2_CD
At-BSK8 / At-FLS2_CD
pGADT7 /At-BSK7
pGADT7 /At-BSK8
At-FLS2_CD$^{D997A}$ / pGBKT7
Sl-BSK7/Sl-FLS2_CD
Sl-FLS2_CD / pGBKT7
pGADT7 / Sl-BSK7
B
-LW
-LWH
$10^{-1}$
$10^{-2}$
i ii iii i ii iii
At-BSK7 / At-BSK7
At-BSK7 / At-BSK8
At-BSK8 / At-BSK8
At-BSK7 / pGBKT7
At-BSK8 / pGBKT7
pGADT7 /At-BSK8
pGADT7 /At-BSK7
Sl-BSK7 / Sl-BSK7
Sl-BSK7 / pGBT9gate
pGADT7 / Sl-BSK7
C
-LW
-LWH
-LWH +10mM3AT
$10^{-1}$
$10^{-2}$
i ii iii i ii iii
Sl-BSK7$_{\Delta1-149\Delta309-486}$/MSP18$_{\Delta sp}$
Sl-BSK7$_{\Delta1-149\Delta309-486}$/pGBKT7
Sl-BSK7$_{\Delta1-149\Delta309-486}$/Sl-FLS2_CD
Sl-BSK7$_{\Delta1-149\Delta309-486}$/pGADT7
Sl-BSK7$_{\Delta150-486}$ /Sl-FLS2_CD
Sl-BSK7$_{\Delta1-308}$ /Sl-FLS2_CD
Sl-BSK7$_{\Delta150-486}$ /pGBT9gate
Sl-BSK7$_{\Delta1-308}$ /pGBT9gate
Sl-FLS2_CD/ pGADT7
Sl-FLS2_CD/pGBKT7

Fig. S7

**Figure S7. Yeast two-hybrid (Y2H) interaction assays reveal BSK–FLS2 association, BSK dimerization, and MSP18–BSK interaction requires the BSK7 middle region.**

(**A**) Y2H assays testing interaction between *A. thaliana* or *S. lycopersicum* BSKs and the cytoplasmic domain (CD) of FLS2, including a kinase-dead FLS2$^{D997A}$ variant (for *A. thaliana*). PJ69-4A yeast strains expressing BSK proteins fused to the GAL4 DNA-binding domain (BD) and FLS2_CD (or FLS2$^{D997A}$) fused to the GAL4 activation domain (AD) were plated on SD medium lacking leucine and tryptophan (-LW), or additionally histidine (-LWH), with or without 3-amino-1,2,4-triazole (3AT) to assess interaction strength. Empty vector controls are included to assess autoactivation.

(**B**) Y2H assays testing homo- and heterodimerization among *A. thaliana* and *S. lycopersicum* BSK7 and BSK8 proteins. Constructs were expressed in yeast with BSKs fused to both BD and AD. Growth on -LWH medium indicates interaction; empty vector controls confirm specificity.

(**C**) Domain mapping of *S. lycopersicum* BSK7 interaction with Mj-MSP18$^{\Delta SP}$ and Sl-FLS2_CD. Yeast expressing truncated Sl-BSK7 variants lacking N-terminal (Δ1–149), C-terminal (Δ309–486), both (Δ1–149Δ309–486), or extended truncations (Δ1–308, Δ150–486) were tested for interaction with Mj-MSP18$^{\Delta SP}$ or Sl-FLS2_CD. Growth on -LWH + 10 mM 3AT medium suggests strong interaction. Results indicate that the middle region of Sl-BSK7 is required for both MSP18 and FLS2 interactions, and that MSP18 binds BSK7 more strongly than FLS2 does.

In all panels, serial dilutions ($10^{-1}$ and $10^{-2}$) of three biological replicates (i, ii, iii) are shown.

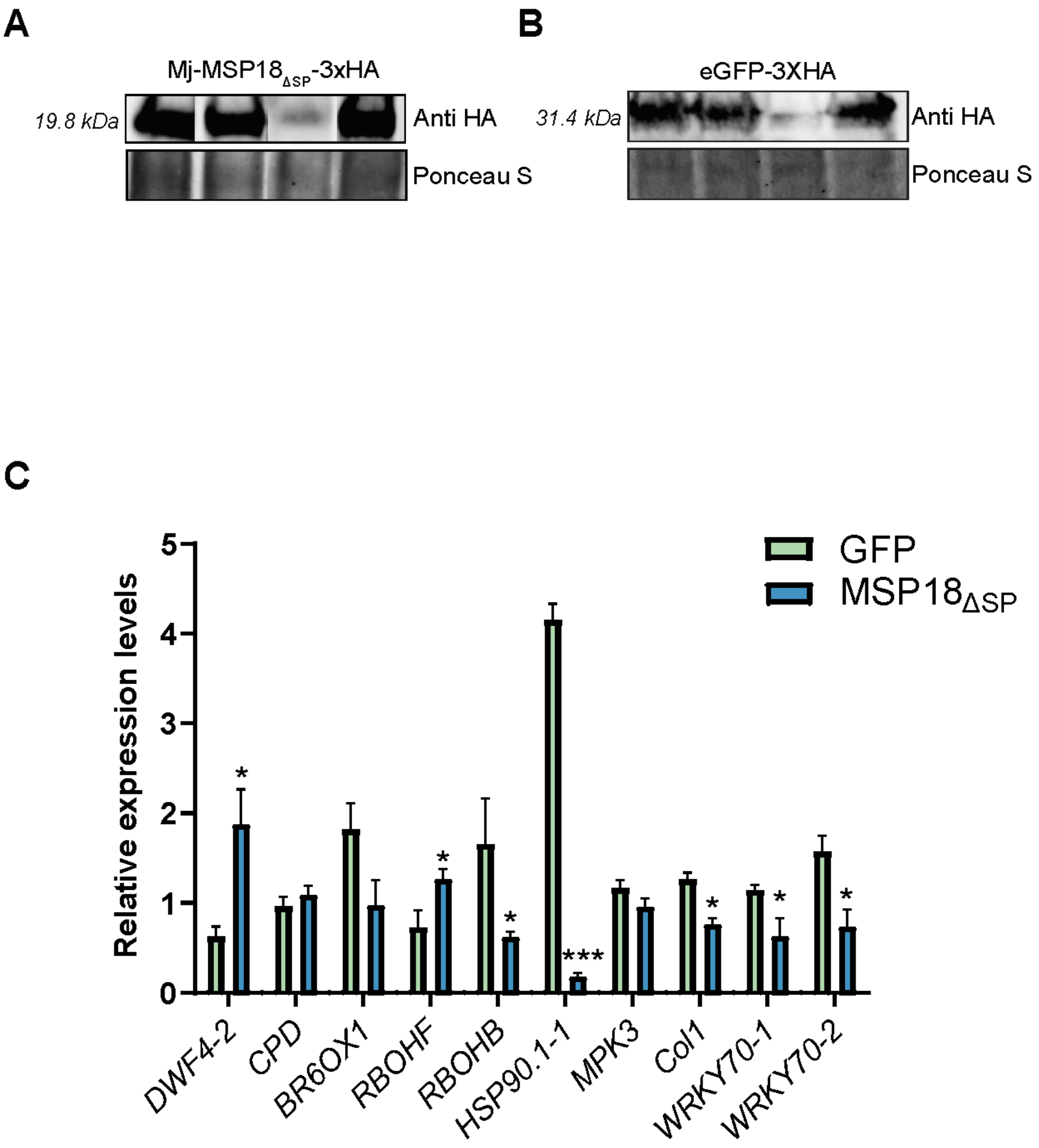
A
Mj-MSP18ΔSP-3xHA
19.8 kDa
Anti HA
Ponceau S
B
eGFP-3XHA
31.4 kDa
Anti HA
Ponceau S
C
GFP
MSP18ΔSP
Relative expression levels
0
1
2
3
4
5
DWF4-2
CPD
BR6OX1
RBOHF
RBOHB
HSP90.1-1
MPK3
Col1
WRKY70-1
WRKY70-2

Fig. S8

**Figure S8. Expression of Mj-MSP18ΔSP and GFP in the and qPCR validation for transcriptional changes caused by Mj-MSP18ΔSP in tomato hairy roots .**

(**A**) Western blot with anti-HA antibody showing the presence of Mj-MSP18$_{\Delta SP}$-3xHA in the tomato hairy roots; the four lanes are from four independently transformed hairy root lines. Expression of the constructs was induced for 24 h with 100 µM estradiol. (**B**) Western blot with anti-HA antibody showing the presence of eGFP-3xHA in the tomato hairy roots; the four lanes are from four independently transformed hairy root lines. Expression of the constructs was induced for 24 h with 100 µM estradiol. (**C**) Validation by RT–qPCR of differently expressed genes in hairy roots upon 24 h of estradiol induction of *Mj-MSP18*$_{\Delta SP}$ compared to *GFP*. Transcript levels were normalized using three reference genes (*ARD, UBE2*, and *DEAD40*), and gene expression using four biological replicates was quantified and expressed relative to *GFP*. Bars represent Std. Error; asterisks indicate significant differences (ns non-significant, * $P < 0.05$, ** $P < 0.01$). qBase plus (Biogazelle, Zwijnaarde, Belgium) software was used to calculate significance and standard error

**Supplemental tables**

**Supplemental table 1.** Identified significantly enriched plant proximal proteins after TurboID **(MSP18 vs GFP)**

**Supplemental Table 2.** Differentially expressed genes (DEG) induced by Mj-MSP18ΔSP compared to eGFP control lines.

**Supplemental Table 3.** Expression profile of randomly selected up- or down-regulated genes in transcriptome data.

**Supplemental Table 4.** Gene ontology (GO) term analysis of significantly downregulated genes in Mj-MSP18ΔSP vs GFP.

**Supplemental Table 5.** Gene ontology (GO) term analysis of significantly upregulated genes in MSP18ΔSP vs GFP.

**Supplemental Table 6.** List of significantly downregulated unique and shared gene between Mj-MSP18$_{\Delta SP}$-induced differentially expressed genes (DEGs) in tomato hairy roots and those downregulated in *Meloidogyne incognita*-infected versus uninfected tomato roots.

**Supplemental Table 7.** Candidate list for Mj-MSP18$_{\Delta SP}$ candidate after Y2H library screen.

**Supplemental Table 8.** The GO term analysis of significant DEGs containing the MYB binding motif (*P* value <0.05).

**Supplemental Table 9.** The GO term analysis of significant DEGs containing the MYC2 binding motif (*P* value <0.05).

**Supplemental Table 10.** List of plasmids used in this study and how they are constructed.

**Supplemental Table 11.** List of all oligos used in this study.

**Supplemental Table 12.** Parameters used for the MaxQuant LFQ search.

**Supplemental Table 13.** Bait sequences provided for MaxQuant analysis.
